# Natural Selection has Shaped Coding and Non-coding Transcription in Primate CD4+ T-cells

**DOI:** 10.1101/083212

**Authors:** Charles G. Danko, Lauren A. Choate, Brooke A. Marks, Edward J. Rice, Zhong Wang, Tinyi Chu, Andre L. Martins, Noah Dukler, Scott A. Coonrod, Elia D. Tait Wojno, John T. Lis, W. Lee Kraus, Adam Siepel

## Abstract

Transcriptional regulatory changes have been shown to contribute to phenotypic differences between species, but many questions remain about how gene expression evolves. Here we report the first comparative study of nascent transcription in primates. We used PRO-seq to map actively transcribing RNA polymerases in resting and activated CD4+ T-cells in multiple human, chimpanzee, and rhesus macaque individuals, with rodents as outgroups. This approach allowed us to measure transcription separately from post-transcriptional processes. We observed general conservation in coding and non-coding transcription, punctuated by numerous differences between species, particularly at distal enhancers and non-coding RNAs. We found evidence that transcription factor binding sites are a primary determinant of transcriptional differences between species, that stabilizing selection maintains gene expression levels despite frequent changes at distal enhancers, and that adaptive substitutions have driven lineage-specific transcription. Finally, we found strong correlations between evolutionary rates and long-range chromatin interactions. These observations clarify the role of primary transcription in regulatory evolution.

---

Following decades of speculation that changes in the regulation of genes could be a potent force in the evolution of form and function^1–3^, investigators have now empirically demonstrated the evolutionary importance of gene regulation across the tree of life^4–12^. Changes in gene expression are primarily driven by mutations to non-coding DNA sequences, particularly those that bind sequence-specific transcription factors^13^. Accordingly, adaptive nucleotide substitutions at transcription factor binding sites (TFBSs)^9,10,14–16^ and gains and losses of TFBSs^17–25^ both appear to make major contributions to the evolution of gene expression. These events are believed to modify a variety of rate-limiting steps early in transcriptional activation^26^. In addition, transcriptional activity is generally correlated with various epigenomic and structural features, including post-translational modifications to core histones, the locations of architectural proteins such as CTCF, and the organization of topological associated domains. Like TFBSs, these features display general conservation across species, yet do exhibit some variation, with differences between species roughly proportional to evolutionary distance^27^. Moreover, differences between species in these features correlate with differences in gene expression^8,24,28–30^.

Nevertheless, many open questions remain about the roles of TFBSs, chromatin organization, and posttranscriptional regulation in the evolution of gene expression. For example, there is a surprisingly limited correlation between differences in binding events and differences in mRNA expression levels^31–33^. Possible reasons for this discordance include non-functional TF binding^31,32,34^, compensatory gains and losses of TFBSs^20,35–38^, difficulties associating distal enhancers with target genes^39^, and a dependency of TF function on chromatin or chromosomal organization^40^. In addition, some changes in mRNA expression appear to be “buffered” at the post-transcriptional level^41–43^. Finally, it remains unclear to what degree epigenomic differences between species are causes and to what degree they are effects of differences in gene expression.

One reason why it has been difficult to disentangle these contributions to gene expression is that expression is typically measured in terms of the abundance of mRNA, which is subject to post-transcriptional processing^44^ and therefore is an indirect measure of the transcription of genes by RNA polymerase II. An alternative and complementary approach is to measure the production of nascent RNAs using Precision Run-On and sequencing (PRO-seq) and related technologies^45–50^. These nascent RNA sequencing methods directly measure active transcription and are highly sensitive to immediate and transient transcriptional responses to stimuli^51^. In addition, they can detect active regulatory elements as well as target genes, because these elements themselves display distinctive patterns of transcription, which are highly underrepresented in RNA-seq data owing to their rapid degradation^34,52,53^. Indeed, the latest nascent RNA sequencing methods, such as PRO-seq^46^, in combination with new computational tools for regulatory element prediction^54^, serve as powerful single-assay systems for both identifying regulatory elements and measuring transcription levels.

With these advantages in mind, we undertook a genome-wide comparative analysis of transcription in primates using PRO-seq. Our comparison of PRO-seq data across species revealed overall conservation in the transcription of both coding and non-coding elements, but also uncovered numerous differences between species. Together, our observations provide new insights into the evolution of transcription in primates.

## Patterns of transcription in resting and activated CD4+ T-cells

We developed nucleotide-resolution maps of RNA polymerase in CD4+ T-cells isolated from five mammalian species. Samples were collected under resting and activated conditions from three unrelated individuals representing each of three primate species—humans, chimpanzees, and rhesus macaques—spanning ∼25-30 million years of evolution (MYR) (Fig. 1a). To compare with studies that focus on longer evolutionary branch lengths, we also collected resting samples from a single individual in each of two rodent species—mouse and rat—which together serve as an outgroup to the primates (∼80 MYR divergence). We used flow cytometry to validate the purity of isolated CD4+ cells (**Supplementary Fig. 1**). In addition, we used measurements of transcriptional activity of T-cell subset markers for T-helper type 1 (Th1), Th2, Th17, T-regulatory, and T-follicular helper cells to demonstrate that the population of CD4+ T-cell subsets within the total CD4+ population was largely similar across species (**Supplementary Fig. 2**).

**Fig. 1.**
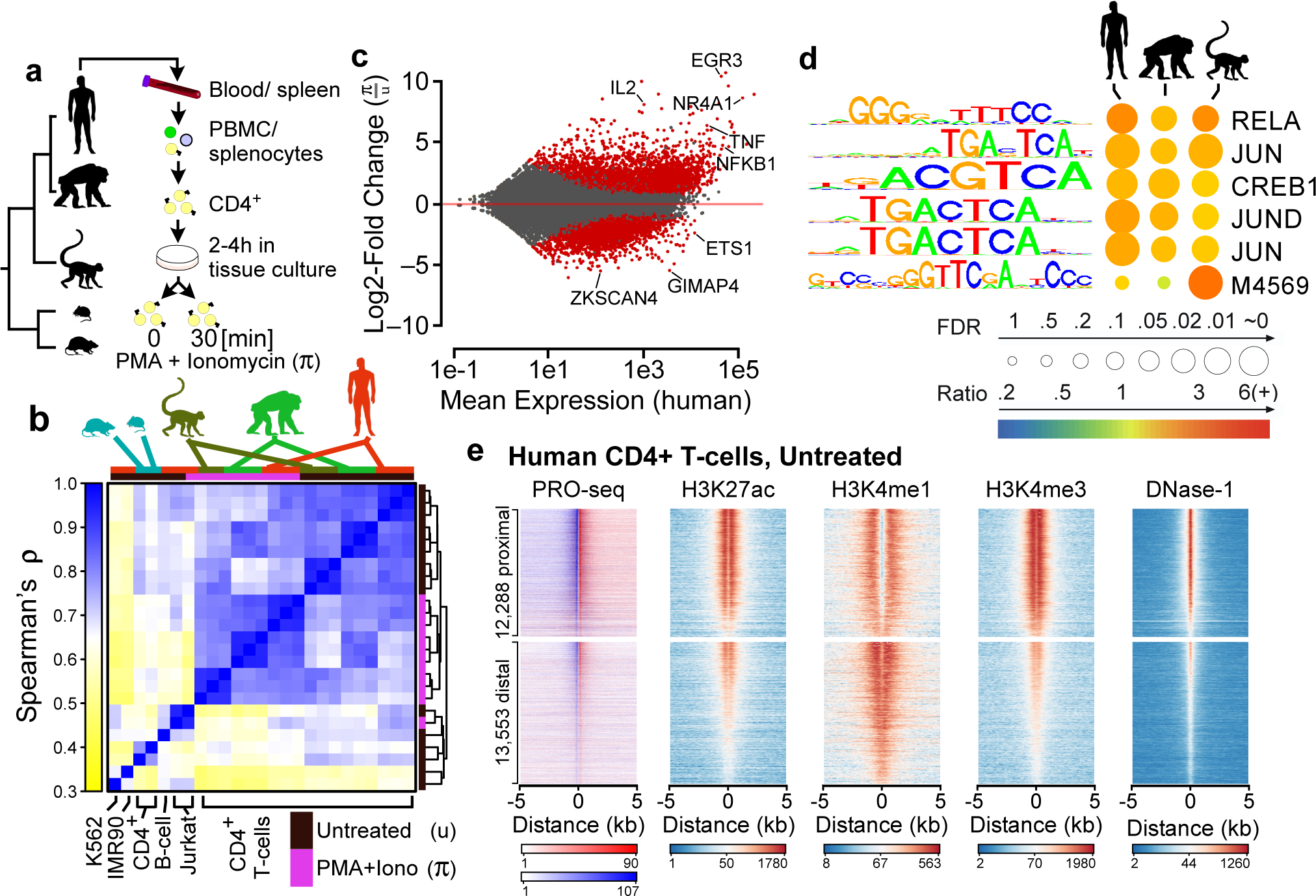
Maps of primary transcription in CD4+ T-cells. **(a)** CD4+ T-cells were isolated from the blood or spleen of individuals from five vertebrate species, including human, chimpanzee, rhesus macaque, mouse, and rat. **(b)** Hierarchical clustering of PRO-seq signal intensities in gene bodies groups CD4+ T-cell samples first by treatment condition and second by species. The color scale represents Spearman’s rank correlation between normalized transcription levels in active gene bodies. Colored boxes (top) represents the species and treatment condition of each sample. **(c)** MA plot shows the log_2_ fold-change following π treatment in human CD4+ T-cells (y-axis) as a function of the mean transcription level in GENCODE annotated genes (x-axis). Red points indicate statistically significant changes (*p* < 0.01). Several classical response genes that undergo well-documented changes in transcript abundance following CD4+ T-cell activation (e.g., *IL2*, *IFNG*, *TNFα*, and *EGR3*) are marked. **(d)** Enrichment of TF binding motifs in TREs that increase transcription levels following π treatment in the indicated species compared to TREs whose transcription abundance does not change. Table shows the Bonferroni corrected p-value, based on a Fisher’s exact test (circle size), and the fold-enrichment over a group of unchanged background sequences (color scale). Motif logos and the candidate transcription factor or Cis-BP motif ID are shown. **(e)** Heatmaps show the distribution of PRO-seq (red and blue indicate transcription on the plus and minus strand, respectively), ChIP-seq for H3K27ac, H3K4me1, and H3K4me3, and DNaseI-seq signal intensity. Plots are centered on transcriptional regulatory elements (TREs) predicted in untreated human CD4+ T-cells using dREG-HD (see Online Methods). All plots are ordered based on the maximum dREG score in the window.

PRO-seq^46,49^ libraries were sequenced to a combined depth of ∼1 billion uniquely mapped reads (∼78-274 million per species) (**Supplementary Table 1**). According to a principal component analysis, the first, second, and third sources of variation in the complete dataset were, respectively, the rodent vs. primate species, variation across the primate species, and the treatment condition (**Supplementary Fig. 3**). Similarly, hierarchical clustering of these data grouped the primate samples first by cell type or treatment condition and subsequently by species, with the rodent samples as outgroups that were more similar to untreated primate CD4+ T-cells than to other samples included in the comparison (Fig. 1b). The correlation between untreated samples decreased linearly with evolutionary time (**Supplementary Fig. 4**), consistent with reports that differences between species arise, on average, at a roughly constant evolutionary rate^27,55^.

To gain further insight into the evolution of the response to CD4+ T-cell stimulation, we compared transcriptional activity under resting and activated conditions within and between species. Here we focused on 42,556 GENCODE-annotated transcription units (TUs) best supported by PRO-seq data for human CD4+ T-cells^56^. In humans, we found that PMA and ionomycin (π) significantly altered the transcription levels of 6,940 (13%) of these TUs (*p* < 0.01, deSeq2^57^; Fig. 1c). Parallel analyses in chimpanzee and rhesus macaque revealed many similarities in transcriptional changes following π treatment (**Supplementary Fig. 5a-b**). We identified a core set of 3,157 TUs that undergo evolutionarily conserved transcriptional changes in all three species following 30 min. of π-treatment, including many of the classical response genes (e.g., IFNG, TNFα, IL2, and IL2RA), as well as numerous novel genes and lincRNAs (**Supplementary Fig. 5a-c**). Active transcriptional regulatory elements (TREs) undergoing changes following π-treatment were enriched for a similar set of transcription factor binding motifs across species, including those for NF-kB and the AP-1 heterodimers FOS and JUN, which are known to be activated by canonical T-cell receptor signaling (Fig. 1d; **Supplementary Note 1**). Thus, the core regulatory principles responsible for T-cell signaling and activation appear to remain broadly conserved across primate evolution.

## Rapid evolutionary changes in transcribed enhancers

We used dREG^54^ to identify 30,357 active TREs in human CD4+ T-cells, based on patterns of enhancer-templated RNA (eRNA) or upstream antisense (uaRNA) transcription evident from PRO-seq data (**Online Methods**). We classified these predicted TREs as either protein-coding promoters or candidate enhancers based on their proximity to gene annotations. The predicted TREs in each group were highly concordant with other marks of regulatory function in human CD4+ T-cells used previously to define groups of candidate enhancers, including acetylation of histone 3 lysine 27 (H3K27ac), mono- and trimethylation of histone 3 lysine 4 (H3K4me1 and H3K4me3), and DNase-I-seq signal^58^ (Fig. 1e). Notably, dREG identified >83% of DNase-I hypersensitive sites (DHS) marked by H3K27 acetylation in human CD4+ T-cells, consistent with prior work suggesting that transcription patterns alone can recover the majority of active enhancers defined by independent criteria^34,52,54^. Furthermore, we identified 88% and 91% of DHSs marked, respectively, by H3K9ac and H4K16ac, two other markers of regulatory function. Taken together, these data suggest that PRO-seq patterns reveal the locations of TREs with high sensitivity. In the analysis that follows, we will refer to these dREG-identified distal TREs as “enhancers” for simplicity.

Extending our dREG analysis to untreated CD4+ T-cells from additional species revealed 71,748 TREs that were active in untreated T-cells in at least one species (ranging between 27,581 and 39,387 TREs in each species). We defined two types of changes between species: (1) changes in the abundance of Pol II at TREs that were present across all species, and (2) complete gains or losses in at least one species (see **Supplementary Note 2, Supplementary Fig. 6,** and **Online Methods**). We found that 52% of enhancers showed evidence of changes in Pol II transcriptional activity in at least one of the three primate species and 81% showed changes at the longer evolutionary distance between primates and rodents (Fig. 2a), similar to recent observations in other systems^10^,24 (**Supplementary Note 3**). Enhancers were predicted to be completely gained or lost at nearly eight times the rate of promoters (35% of enhancers; 12% of promoters; *p* < 2.2e-16 by Fisher’s exact test), consistent with recent observations based on H3K27ac and H3K4me3^24^. By contrast, TREs induced by π treatment were much more likely to be conserved, and showed similar conservation at promoters and enhancers (Fig. 2a).

**Fig. 2.**
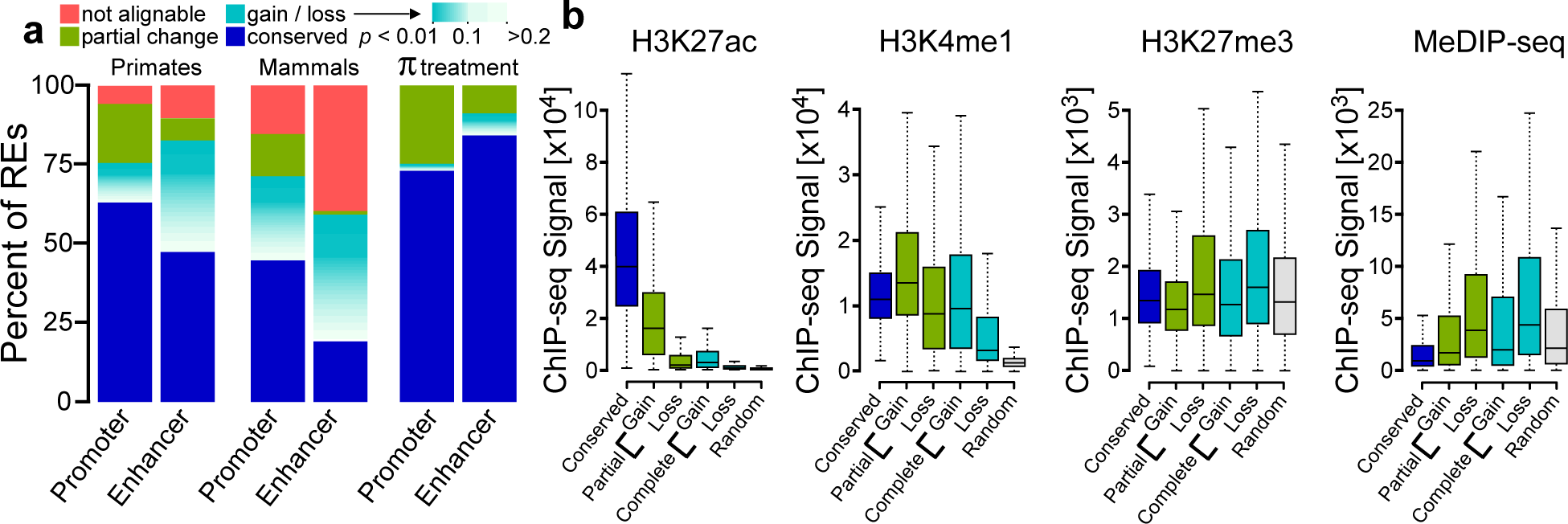
Frequency of changes in TRE transcription. **(a)** The fractions of TREs active in untreated CD4+ T-cells that are present in the human reference genome and are conserved across all species (blue), are not detectable and are therefore inferred as gains or losses (teal-white) or undergo significant changes (green) in at least one species, or fall in regions for which no ortholog occurs in at least one of the indicated genomes (pink). Inferred gains or losses are colored according to the FDR corrected p-value associated with changes in RNA polymerase abundance (deSeq2). Plots labeled “Primate” illustrate frequency of changes in a three-way comparison of human, chimpanzee, and rhesus macaque focusing on the untreated condition, whereas those labeled “Mammal” summarize a five-way comparison also including rat and mouse. π treatment denotes a comparison between human untreated and PMA+Ionomycin treated CD4+ T-cell samples. **(b)** Boxplots show the ChIP-seq signal near dREG sites classified as conserved, gains, losses, or complete losses for the indicated chromatin or DNA modification in units of reads per kilobase. The box represents the 25th and 75th percentile. Whiskers represent 1.5 times the interquartile range, and points outside of this range are not shown.

Next we tested whether evolutionary changes in transcriptional activity correlate with the enrichment of other marks of active enhancers. Predicted lineage-specific human enhancers were enriched for both active and repressive enhancer marks (Fig. 2b; **Supplementary Fig. 7**). Whereas apparent human gains were enriched for high levels of the active enhancer marker H3K27ac, sites with reduced transcriptional activity in humans showed much lower enrichments of H3K27ac. Furthermore, locations at which the dREG signal was completely lost in a human-specific fashion displayed levels of H3K27ac approaching those of randomly selected background sites (Fig. 2b). Intriguingly, many of the losses on the human branch retained H3K4me1, which marks both active and inactive enhancers^59,60^, and these losses displayed higher levels of chromatin marks associated with transcriptional repression than a random background (Fig. 2b), indicating that, at least in some cases, an active ancestral primate enhancer retains a ‘poised’ chromatin state in human, despite losing both transcriptional activity and H3K27ac. Thus, evolutionary changes in poised and active marks may commonly occur as distinct events.

## Transcriptional changes correlate with DNA sequence differences

To investigate whether changes in TRE activity are accompanied by changes in DNA sequence, we compared phyloP sequence conservation scores^61^ at transcriptionally conserved TREs with phyloP scores at TREs that display evolutionary changes in transcription. Because signatures of sequence conservation in TREs are likely to be most pronounced in transcription factors binding sites (TFBS), we restricted our sequence conservation analyses to matches to 567 clusters of TF binding motifs selected based on their distinct DNA binding specificities (see **Online Methods**)^62,63^. We required that motif matches were present in dREG sites, and adopted a threshold at which nearly half of the motifs discovered were bona fide TFBSs, as measured by ChIP-seq (positive predictive value [PPV]= 0.47).

TFBSs found in transcriptionally conserved dREG sites showed a marked enrichment for higher phyloP scores relative to surrounding regions, indicating local sequence conservation (Fig. 3a). By contrast, TFBSs in lineage-specific dREG sites had substantially reduced enrichments in phyloP scores (Fig. 3a, **cyan/blue**). Notably, TFBSs in dREG sites lost on the human lineage showed enhanced conservation compared with those in human-specific gains. This observation is consistent with losses evolving under conservation in other mammalian species (which contribute to the phyloP scores) and gains emerging relatively recently. Each of these patterns was robust to corrections for potentially confounding differences in the distribution of sites, as well to choices of motif score thresholds (**Supplementary Fig. 8a**). Relaxing the motif score threshold to provide sensitivity for larger numbers of TFBSs at the expense of specificity, revealed patterns of conservation that correlate with the information content of positions within the DNA sequence motif (**Supplementary Fig. 8b**), further supporting TF binding as the functional property driving sequence conservation at these sites. Together, these analyses support the hypothesis that the sequences in TFBSs are a primary driver of transcriptional differences between species.

**Fig. 3.**
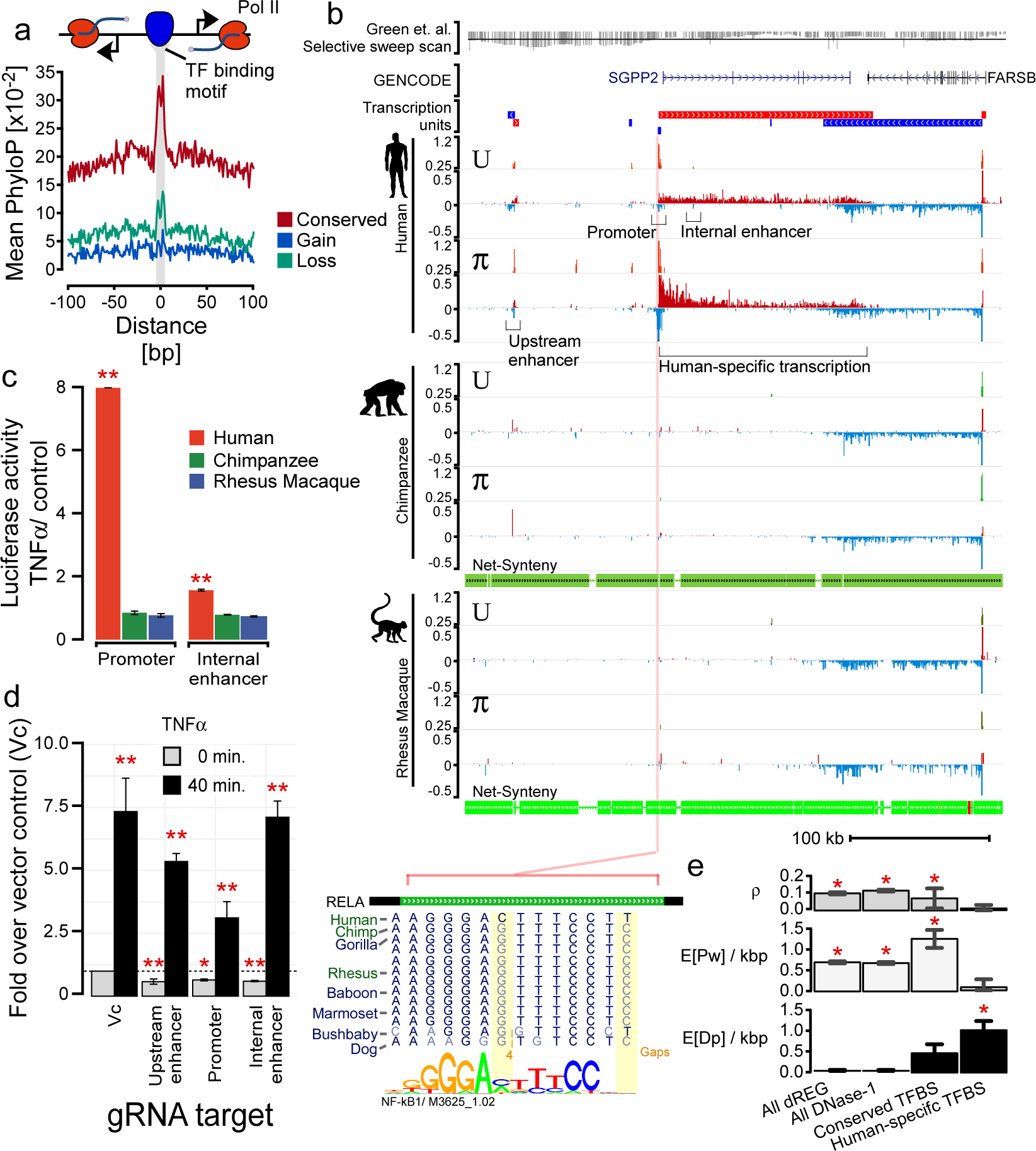
Evolutionary changes in TRE transcription correlate with DNA sequence conservation. **(a)** Mean phyloP scores near TFBSs that are conserved (red), gained (blue), or lost (cyan) on the human branch. Motifs (score > 10) are at least 100 bp from the nearest annotated exon. **(b)** UCSC Genome Browser track shows transcription near *SGPP2* and *FARSB* in untreated (U) and PMA+ionomycin (π) treated CD4+ T-cells isolated from the indicated primate species. PRO-seq tracks show transcription on the plus (red) and minus (blue) strands. Axes for the PRO-seq data are in units of reads per kilobase per million mapped (RPKM). Transcription units inferred from the PRO-seq data are shown above the plot. dREG tracks show the distribution of dREG signal. The Green et. al. (ref^64^) selective sweep scan track (top) represents the enrichment of derived alleles in modern human where Neanderthal has the ancestral allele. Points below the line represent a statistically significant number of derived alleles in modern human (line indicates a Z-score of -2). Net synteny tracks show the position of regions that have one-to-one orthologs in the chimpanzee and rhesus macaque genomes. **(c)** Luciferase signal driven by the *SGPP2* promoter or the internal enhancer in MCF-7 cells using DNA from each primate species. Bars show the mean fold-induction following 3 hours of stimulation with TNFα. Error bars represent the standard error of the mean. Red ** denotes *p* < 1e-3 by a two-tailed t-test. **(d)** Transcription of *SGPP2* using primers targeting intron 1 following 0 or 40 min. of TNFα treatment after silencing the indicated TRE using dCAS9-KRAB. Bars represent the median of three independent biological replicates of two gRNAs targeting the promoter, three targeting the internal enhancer, and four targeting the upstream enhancer. Error bars represent the standard error. Red **\*** denotes *p* < 5e-2 and **\*\*** *p* < 5e-3 by a two-tailed t-test. **(e)** INSIGHT estimates of the fraction of nucleotides under selection (ρ), the expected density of segregating polymorphisms under weak negative selection (E[Pw]/kbp), or the expected density of human nucleotide substitutions driven by positive selection (E[Dp]/kbp) in human populations in the indicated class of sites. Red **\*** denotes conditions significantly enriched over random background sequences (p < 0.01; two-tailed ?^2^-test).

We searched for examples of DNA sequence differences that might be responsible for transcriptional changes following π treatment, hypothesizing that they might be characterized more easily than sequences responsible for transcription in the untreated condition, because these transcriptional changes were likely driven by a limited number of master TFs^64^ (Fig. 1e). In one example, we found nucleotide substitutions in three apparent NF-kB binding sites in the proximal promoter and an internal enhancer of *SGPP2* that correlate with differences in *SGPP2* expression **(Fig. 3b; Supplementary Fig. 9**). Two of these putative binding sites were bound by NF-kB in human cell lines according to ChIP-seq data from ENCODE. Moreover, substitutions observed in human were found to disrupt the same position in the motif as NF-kB binding QTLs^65^ (see **Online Methods**), and showed a general trend toward higher NF-kB binding in the human alleles **(Supplementary Fig. 9**). To test the hypothesis that observed DNA sequence changes produced differential transcriptional activity, we cloned DNA from each primate species into a reporter vector driving luciferase activity in an MCF-7 cell model, which recapitulates the primary transcriptional features of the *SGPP2* locus^66^ (**Supplementary Fig. 9**). Differences in basal luciferase activity were generally concordant with those observed between species **(Supplementary Fig. 10**). Moreover, both the proximal promoter of *SGPP2* and the internal enhancer both activated luciferase expression more strongly following NF-kB activation when human DNA was cloned, but not with orthologous DNA from the other primates (Fig. 3c).

To determine whether these TREs affect the expression of *SGPP2* in its native genomic context, we silenced each TRE by using CRISPRi, which targets a catalytically dead CAS9 fused to the Krüppel-associated box repressor (dCAS9-KRAB), to specifically tri-methylate lysine 9 of histone 3 (H3K9me3)^67^. Three independent single-guide RNAs (sgRNAs) targeting the internal enhancer and two designed for the proximal promoter reduced *SGPP2* transcription to 50-60% of its resting level (*p* = 1.5e-3 and 2.6e-2, respectively, by a two-tailed t-test; Fig. 3d), consistent with these TREs directly contributing to *SGPP2* expression in MCF-7 cells. Three sgRNAs targeting the upstream enhancer also had a significant effect on *SGPP2* expression (*p* = 1.8e-4). Notably, the genome assemblies for chimpanzee and rhesus macaque harbor deletions of this upstream TRE that appear likely to affect its activity (**Supplementary Fig. 9**). However, although silencing individual enhancers reduced the transcription level of *SGPP2* following NF-kB activation, silencing individual enhancers was insufficient to completely abolish induction of *SGPP2* (Fig. 3d). Taken together, our findings show that at least two of the three TREs regulating *SGPP2* drove expression patterns matching PRO-seq data in a reporter assay, but none completely explained *SGPP2* activation *in situ*. These observations suggest that that multiple causal substitutions in NF-kB binding sites may work in concert to achieve human-specific activation of *SGPP2*.

In several cases, as with *SGPP2*, we observed numerous nucleotide substitutions within individual or clustered TFBSs. These clusters of substitutions are highly unlikely to occur by chance and suggest that positive selection may have driven adaptation of these binding sites. Indeed, *SGPP2* falls in a region recently identified as having an excess of derived alleles in modern humans compared with the sequenced Neanderthal (Fig. 3d)^68^, potentially consistent with recent positive selection driving evolutionary changes in *SGPP2* transcription. To more directly measure the impact of positive selection, we used INSIGHT^69^ to compare patterns of within-species polymorphism and between-species sequence divergence in TREs that had undergone human lineage-specific transcriptional changes. This analysis indicated that dREG sites are most strongly influenced by weak negative selection (Fig. 3f), based on an excess of low-frequency derived alleles in human populations, as has been reported previously for regulatory sequences^9^. Nevertheless, TREs with lineage-specific transcriptional changes in human CD4+ T-cells showed reduced weak negative selection and were strikingly enriched for adaptive nucleotide substitutions (*p* < 0.01 INSIGHT likelihood ratio test; Fig. 3f), consistent with positive selection at these sites. We estimate a total of at least 121 adaptive substitutions since the human/chimpanzee divergence within TFBSs that undergo transcriptional changes in human CD4+ T-cells. Despite limited power to detect the specific contributions of many individual TFs at our stringent motif match score threshold, we did note significant excesses of putatively adaptive substitutions in the predicted binding sites of several TFs, including motifs recognized by forkhead box family, POU-domain containing, and ELF/ETS family (**Supplementary Fig. 11**; *p* < 0.01, INSIGHT likelihood ratio test). These estimates highlight the substantial contribution of adaptive evolutionary changes in TFBSs that may influence the transcriptional activity of TREs.

## Correlation between protein-coding and non-coding transcription

We noticed that evolutionary changes in protein-coding gene transcription frequently correlate with changes in non-coding transcription units (TU) located nearby. To examine this pattern more generally, we adapted our recently reported hidden Markov model (HMM)^70^ to estimate the location of TUs genome-wide, based on patterns of aligned PRO-seq reads and the location of TREs. Using this method, we annotated 54,793 TUs active in CD4+ T-cells of at least one of the primate species, approximately half of which overlap existing GENCODE annotations or their associated upstream antisense RNAs (**Supplementary Fig. 12a**). A cross-species comparison of the transcription levels for various TU classes (Fig. 4a) revealed that non-coding RNAs evolve in expression most rapidly and protein-coding genes evolve most slowly. GENCODE-annotated lincRNAs undergo evolutionary changes in expression about as frequently as the unannotated non-coding RNAs predicted by our HMM, which are likely enriched for bi-directionally unstable eRNA species. The broad similarity in evolutionary conservation between these two non-coding RNA classes may be consistent with observations that some lincRNAs function as enhancers for nearby protein-coding genes, and that stable accumulation of the transcript is dispensable for this biological function^71^.

**Fig. 4.**
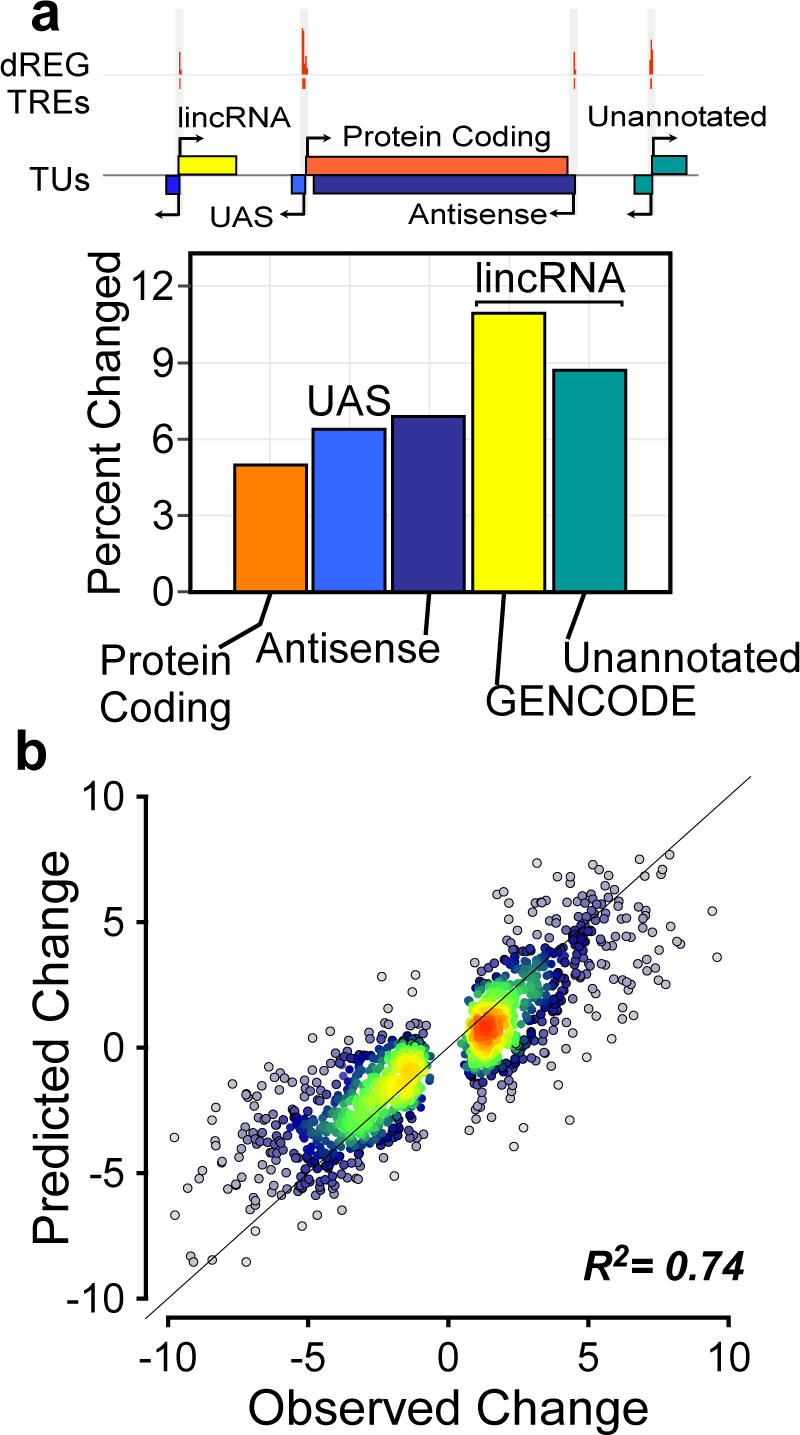
Changes in non-coding RNA transcription predict changes in gene transcription. **(a)** The fraction of each indicated class of RNAs that undergo changes in transcription in human CD4+ T-cells (see Online Methods). The relationships among the indicated classes of transcription units are depicted at top. **(b)** Scatterplot shows the magnitude of changes in transcription predicted for protein-coding genes using changes in the transcription of nearby non-coding RNAs (y-axis) as a function of changes observed (x-axis). The line has a slope of 1 and an intercept of 0.

We next measured the extent to which non-coding and protein-coding transcriptional activities are correlated through evolutionary time. We found that evolutionary changes in protein-coding gene expression among any of the primate species were highly correlated with those at both upstream (Pearson’s *R* = 0.85, *p* < 2.2e-16) and internal ( *R* = 0.66, *p* < 2.2e-16) antisense transcripts of the same genes. Moreover, changes in the transcriptional activity of gene promoters correlate with changes in the activity of matched distal enhancers defined by various criteria: namely, enhancers to which the promoters loop according to cell-type matched ChIA-PET data ( *R* = 0.45-0.61, p < 2.2e-16; depending on analysis assumptions)^72^, enhancers located nearby the promoters *(R* = 0.69, p < 2.2e-16), or enhancers that share the same topological associated domain as the promoter ( *R* = 0.62, p < 2.2e-16)^73^. Using a generalized linear model to integrate expression changes in multiple types of TUs, we can explain 74% of the variance in gene transcription levels when we observe differences between species ( *R*^2^ = 0.74 in a held-out set of sites, *p* < 2.2e-16; Fig. 4b) based on the activities of looped TREs, nearby TREs in the same topological associated domain, internal antisense TUs, and the upstream antisense TU. Thus evolutionary changes that result in differences in Pol II recruitment to protein-coding genes are well correlated across all interacting TREs, indicating a shared evolutionary pressure at proximal and distal TREs.

## Rates of Enhancer Evolution Vary with Evidence for Gene Interactions

Despite this overall positive correlation, transcription at enhancers evolves rapidly and is frequently unaccompanied by transcriptional changes at nearby protein-coding genes. For example, *CCR7* transcription is highly conserved among both primate and rodent species (Fig. 5a; **Supplementary Fig. 2**) in spite of several apparent changes in enhancer activity within the same locus (gray vertical bars). These findings are consistent with recent observations that changes in enhancers within densely populated loci often do not have appreciable effects on the transcription of genes within the locus^36,37^.

**Fig. 5.**
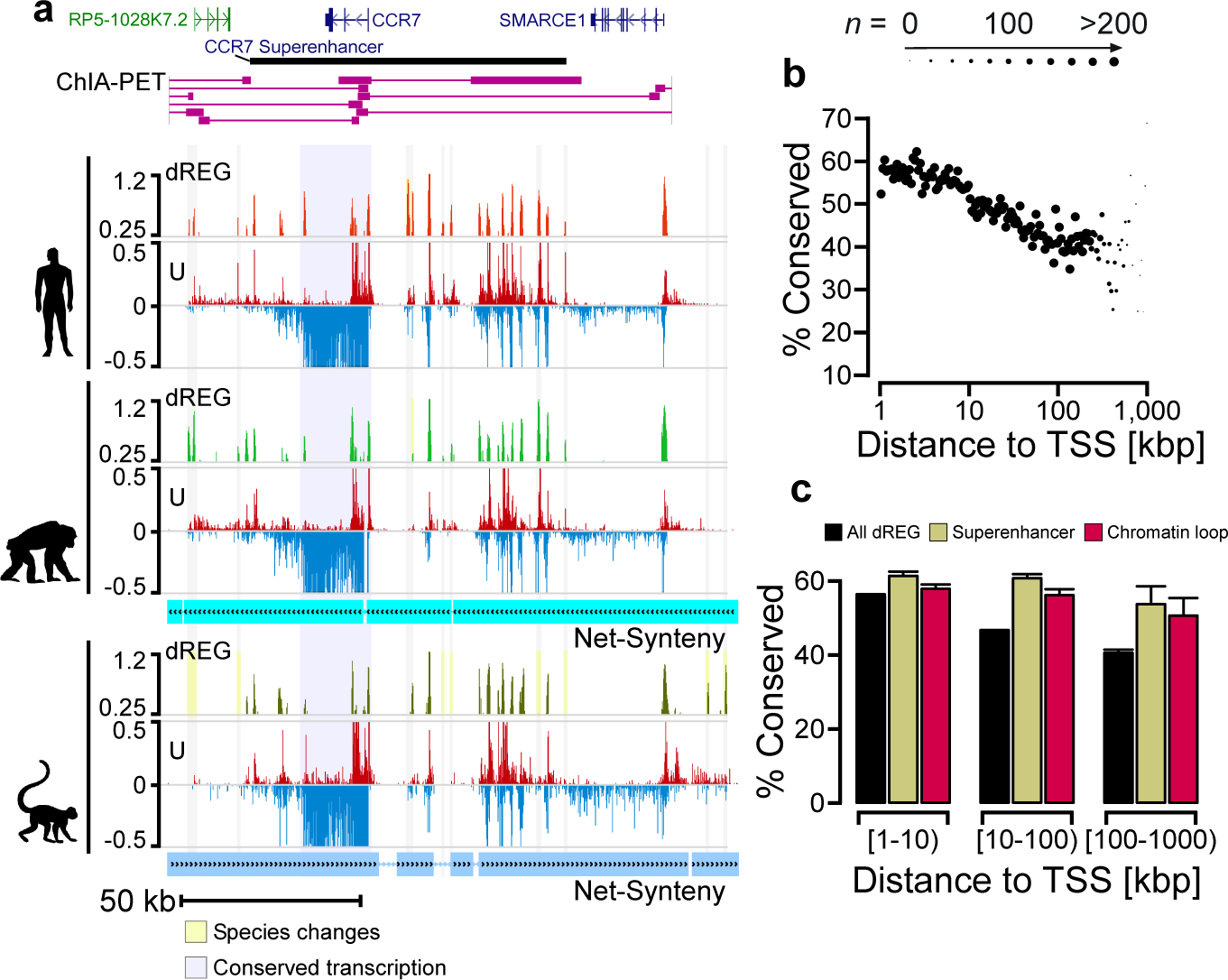
TRE conservation correlates with loop interactions and distance to gene promoters. **(a)** UCSC Genome Browser tracks show transcription, dREG signal, and ChIA-PET loop interactions near the *CCR7* superenhancer in the human genome. PRO-seq tracks show transcription on the plus (red) and minus (blue) strands in units of RPKM. Net synteny tracks show regions of one-toone orthology with the chimpanzee and rhesus macaque genomes. **(b)** Scatterplot shows the percentage of TREs conserved among all three primate species (y-axis) as a function of distance, either upstream or downstream, from the nearest annotated protein-coding transcription start site (x-axis). The size of each point represents the amount of data in the corresponding distance bin. The percentage of all dREG sites that are conserved in each indicated class of TRE. TREs are separated into three bins based on the distance relative to the nearest transcription start site. Error bars reflect a 1,000-sample bootstrap.

To explain this effect, we searched for genomic features correlated with conservation of transcription at enhancers, focusing on untreated CD4+ T-cells in order to leverage the large amount of public data available for this cell type. Not surprisingly, one of the features most strongly correlated with transcriptional conservation at enhancers is the distance from the nearest transcription start site of a protein-coding gene (Fig. 5b). In particular, more than half of enhancers located within 10 kbp of an annotated TSS are shared across all three primate species, whereas for distal enhancers located between 100 kbp to 1 Mbp from a TSS that fraction drops to roughly a third. This relationship is driven by lineage-specific gains or losses of enhancer activity, and to a lesser extent by changes in TRE activity levels, rather than by differences in the alignability of orthologous DNA (**Supplementary Fig. 13**).

These simple distance-based observations, however, ignore the critical issue of chromatin interactions between enhancers and promoters. To account for such loop interactions, we extracted 6,520 putative TRE interactions from Chromatin Interaction Analysis with Paired End Tag sequencing (ChIA-PET) data recognizing loops marked with H3K4me2 in human CD4+ T-cells^72^. We found that 55% of enhancers that participate in these loops were conserved between primate species compared to only 47% of non-looped enhancers (Fig. 5c; *p* = 5.6e-4, Fisher’s exact test). Moreover, higher transcriptional conservation at looped enhancers does not depend on the distance to the transcription start site. Parallel analysis of promoter-capture Hi-C data^74^ revealed that the strength of chromatin interaction was correlated with evolutionary conservation of distal TREs, corroborating the result obtained using ChIA-PET (*p* < 1e-3, bootstrap test). We observed similar levels of conservation at recently defined super-enhancers^75^, although this conservation may simply reflect an enrichment for loop interactions (48% of TREs in super-enhancers loop according to ChIA-PET, compared to 15% of all TREs). Looped enhancers were also enriched for elevated phyloP scores relative to either non-looped enhancers or randomly selected DNA sequences ( **Supplementary Fig. 14;** phyloP > 0.75; *p* < 2.2e-16, Wilcoxon Rank Sum Test). That the enhancers participating in loop interactions are more highly conserved at both the transcription and DNA-sequence levels indicates that these enhancers have a disproportionately large effect on fitness, presumably owing to a more direct role in transcriptional regulation.

## Enhancer-Promoter Interactions Contribute to Constraint on Gene Transcription Levels

Distal loop interactions do not fully account for the disparity between enhancers and promoters in evolutionary rates—even looped enhancers still evolve significantly faster than promoters (*p* = 3e-3, Fisher’s exact test). We hypothesized that redundancy in enhancers may help to explain rapid enhancer evolution. More specifically, we asked whether redundancy makes protein-coding genes regulated by multiple distal TREs, such as *CCR7* (Fig. 5a), more robust to enhancer turnover than those influenced by fewer distal TREs. Indeed, we found that evolutionary conservation of promoter TRE transcription is remarkably strongly correlated with the number of loop interactions a promoter has with distal sites (Fig. 6a, weighted Pearson’s correlation = 0.87; *p* < 1e-3 by a bootstrap test). A similar trend was observed between the number of loop interactions made by a target promoter and DNA sequence conservation in transcription factor binding motifs at the promoter, although the effect was weaker and did not meet our criteria for statistical significance (Fig. 6b, weighted Pearson’s correlation = 0.65; *p* = 0.07 by a bootstrap test).

**Fig. 6.**
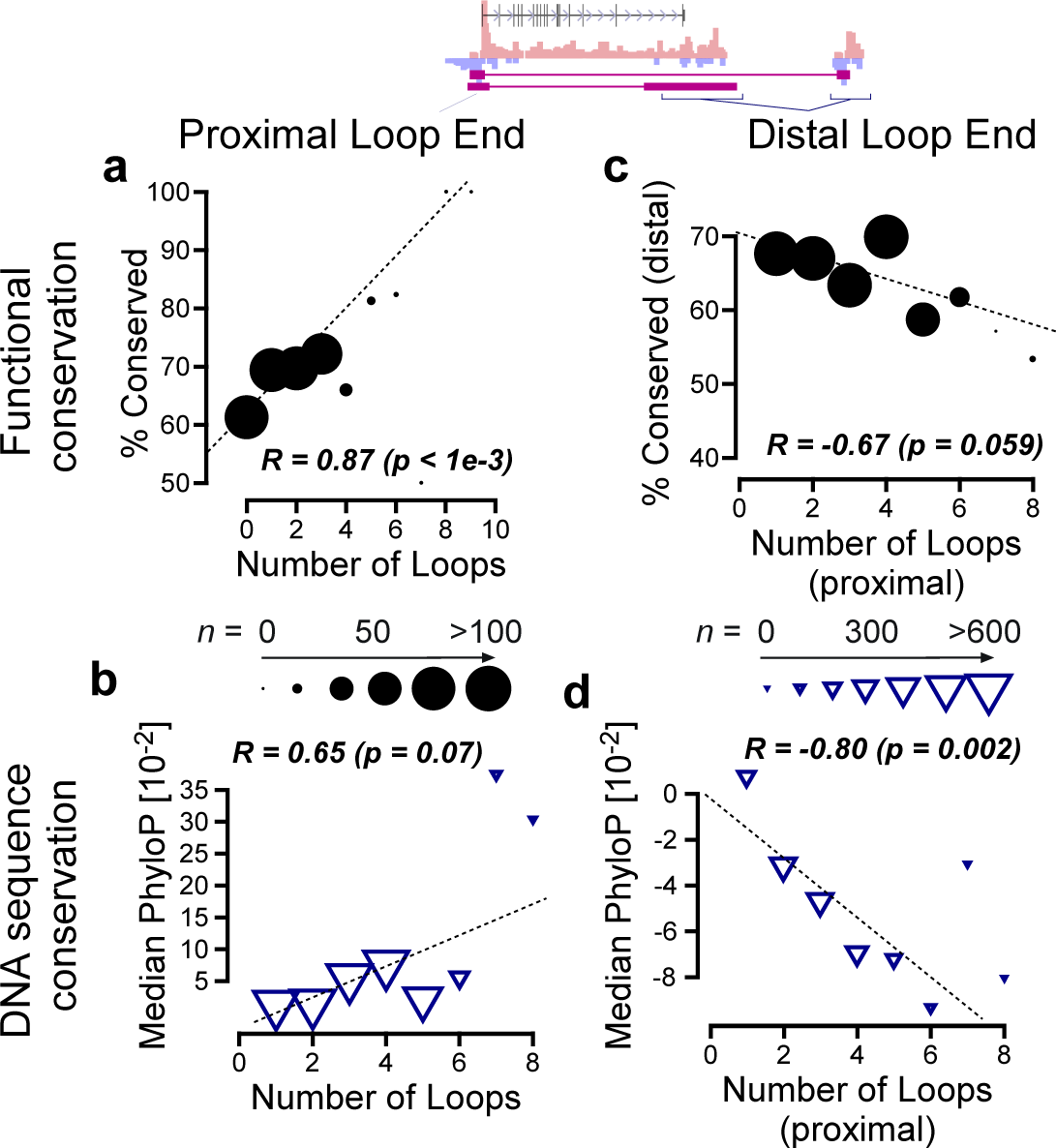
Stabilizing selection on protein coding gene transcription. **(a-b)** Scatterplot shows promoter conservation **(a)** or DNA sequence conservation **(b)** as a function of the number of loop interactions made by that site to distal sites across the genome (x-axis). **(c-d)** TRE conservation **(c)** or DNA sequence conservation **(d)** as a function of the number of loop interactions made by the sequence at the distal end of the loop interaction (x-axis). In all panels the size of each point is proportional to the number of examples in the corresponding bin, following the scale shown in the center.

But how does redundancy in distal TREs relate to the evolutionary conservation of the distal TREs themselves? If redundant distal TREs compensate for one another in some way, perhaps each one will be less, rather than more, conserved when their associated promoters have larger numbers of loop interactions. To address this question, we examined the rate of conservation of looped distal TREs as a function of the number of loops in which their gene-proximal partners participated. We found that DNA sequence conservation in putative TFBSs negatively correlates with the number of loops at the proximal end (Fig. 6d; weighted Pearson’s correlation = –0.80; *p* = 2e-3 by a bootstrap test). We noted a similar trend toward a negative correlation between the conservation of distal TRE transcription and the number of loop interactions (weighted Pearson’s correlation = –0.67; *p* = 0.059, two-sided bootstrap test, Fig. 6c). These results suggest that each associated distal TFBS is individually less essential at genes having larger numbers of loop interactions with distal sites, and they are therefore consistent with a model in which such TFBSs are more freely gained and lost during evolution. Taken together, our results imply that distance, looping, and redundancy of enhancers all contribute to constraints on the evolutionary rates of changes in gene transcription.

## Discussion

We have carried out the first comparative analysis of primary transcription in any phylogenetic group, focusing on CD4+ T-cells in primates. Using PRO-seq and various computational tools, we estimated the locations and abundance of transcription units with high resolution and accuracy. In comparison to previous studies in primates^29,33,76–78^, this approach separated primary transcription from post-transcriptional processing, allowing us to study eRNAs, lincRNAs, and other rapidly degraded non-coding RNAs, as well as protein-coding genes. We found clear relationships between the DNA sequences of TFBSs and differential transcription across species and treatment conditions. We also found evidence that some transcriptional changes in humans were driven by adaptive evolution in nearby binding sites. Overall, our study provides new insights into the mode and tempo of recent evolutionary changes in transcription in primates.

Perhaps our most striking observation is that many non-coding transcription units, particularly eRNAs and lincRNAs, have undergone rapid evolutionary changes in comparison to protein-coding genes. Similar observations have been reported previously for lincRNAs^79^, but, to our knowledge, the observation for eRNAs is new, and it raises a number of questions. First, why are some enhancers more conserved than others? In particular, we find that enhancers proximal to, or that loop to, annotated promoters tend to be most constrained (**Fig 5b-c**). These enhancers may simply be most crucial for activating their target genes, but other factors may also contribute to their constraint. For example, perhaps these enhancers are enriched for tissue-specific functions, and are less constrained due to reduced pleiotropy^80^. Or perhaps many of them simply are not functional at all, and are transcribed as a by-product of other processes.

Second, how do protein-coding genes maintain stable transcription levels across species despite the rapid turnover of associated enhancers? One possibility is that many rapidly evolving enhancers are either not functional or act on targets other than the ones we have identified. However, several of our findings argue against this possibility; for example, we find that even looped enhancers, for which we have direct evidence of a promoter interaction, evolve significantly faster than promoters, and that eRNA conservation is strongly correlated with the number of loop interactions at associated promoters (Fig. 6). An alternative explanation, which appears more plausible to us, is that stabilizing selection on transcription levels drives enhancers to compensate for one another as they undergo evolutionary flux. This explanation would be compatible with reports from model systems^36,38^. Our finding that sequence conservation at distal enhancers is negatively correlated with the number of loop interactions at associated promoters **(Supplementary Fig. 14**) is also consistent with this explanation. The possibility of pervasive stabilizing selection on transcription levels in primates has been noted previously based on RNA-seq data^81^, but our data allow for more direct observations of both active transcription and associated regulatory elements.

A third question is, if most transcribed enhancers do indeed influence gene expression, then why are so many of them weakly maintained by natural selection? At present, we can only speculate on the answer to this question. One possibility is that some of the apparent turnover events we have observed actually represent enhancers that have simply switched cell types in activity, as has been reported in some cases^19^. But it is also possible that selection tends to act diffusely on enhancers across an entire locus, rather than strongly on individual enhancers, as has been proposed in cancer evolution^82^. Our observation that multiple DNA sequence changes at the *SGPP2* locus appear functional provides some initial support this this hypothesis. It will be possible to evaluate these hypotheses more rigorously as better data describing enhancers and enhancer-promoter interactions across many cell types become available for these and other groups of species.

## Methods

### Multiple species PRO-seq library generation

#### Isolation of primate CD4+ T-cells

All human and animal experiments were done in compliance with Cornell University IRB and IACUC guidelines. We obtained peripheral blood samples (60-80 mL) from healthy adult male humans, chimpanzees, and rhesus macaques. Informed consent was obtained from all human subjects. To account for within-species variation in gene transcription we used three individuals to represent each primate species. Blood was collected into purple top EDTA tubes. Human samples were maintained overnight at 4C to mimic shipping non-human primate blood samples. Blood was mixed 50:50 with phosphate buffered saline (PBS). Peripheral blood mononuclear cells (PBMCs) were isolated by centrifugation (750x g) of 35 mL of blood:PBS over 15 mL Ficoll-Paque for 30 minutes at 20C. Cells were washed three times in ice cold PBS. CD4+ T-cells were isolated using CD4 microbeads (Miltenyi Biotech, 130-045-101 [human and chimp], 130-091-102 [rhesus macaque]). Up to 10^8^ PBMCs were resuspended in binding buffer (PBS with 0.5% BSA and 2mM EDTA). Cells were bound to CD4 microbeads (20uL of microbeads/10^7^ cells) for 15 minutes at 4C in the dark. Cells were washed with 1-2 mL of PBS/BSA solution, resuspended in 500uL of binding buffer, and passed over a MACS LS column (Miltenyi Biotech, 130-042-401) on a neodymium magnet. The MACS LS column was washed three times with 2mL PBS/BSA solution, before being eluted off the neodymium magnet. Cells were counted in a hemocytometer.

#### Isolation of CD4+ T-cells from mouse and rat

Spleen samples were collected from one male mouse (FVB) and one male rat (Albino Oxford) that had been sacrificed for IACUC-approved research not related to the present study. Dissected spleen was mashed through a cell strainer using a sterile glass pestle and suspended in 20 mL RPMI-1640. Cells were pelleted at 800xg for 3 minutes and resuspended in 1-5mL of ACK lysis buffer for 10 minutes at room temperature to lyse red blood cells. RPMI-1640 was added to a final volume 10 times that used for ACK lysis (10-40 mL). Cells were pelleted at 800xg for 3 minutes, counted in a hemocytometer, and resuspended in RPMI-1640 to a final concentration of 250,000 cells per ml. CD4+ T-cells were isolated from splenocytes using products specific for mouse and rat (Miltenyi Biotech, 130-104-453 [mouse], 130-090-319 [rat]) following instructions from Miltenyi Biotech, and as described above.

#### T-cell treatment and PRO-seq library generation

CD4+ T-cells were allowed to equilibrate in RPMI-1640 supplemented with 10% FBS for 2-4 hours before starting experiments. Primate CD4+ T-cells were stimulated with 25ng/mL PMA and 1mM Ionomycin (P/I or π) or vehicle control (2.5uL EtOH and 1.66uL DMSO in 10mL of culture media). We selected the minimum concentrations which saturate the production of IL2 and IFNG mRNA after 3 hours of treatment (data not shown). A 30 min. treatment duration was selected after observing a sharp increase in ChIP-qPCR signal for RNA Pol II phosphorylated at serine 5 on the C-terminal domain on the IFNG promoter at 30 min. (data not shown). To isolate nuclei, we resuspended cells in 1 mL lysis buffer (10 mM Tris-Cl, pH 8, 300 mM sucrose, 10 mM NaCl, 2 mM MgAc2, 3 mM CaCl2 and 0.1% NP-40). Nuclei were washed in 10 mL of wash buffer (10 mM Tris-Cl, pH 8, 300 mM sucrose, 10 mM NaCl and 2 mM MgAc2) to dilute free NTPs. Nuclei were washed in 1 mL, and subsequently resuspended in 50 μL, of storage buffer (50 mL Tris-Cl, pH 8.3, 40% glycerol, 5 mM MgCl2 and 0.1 mM EDTA), snap frozen in liquid nitrogen and kept for up to 6 months before making PRO-seq libraries. PRO-seq libraries were created exactly as described previously^46^. In most cases, we completed library preps with one member of each species (usually one human, chimpanzee, and rhesus macaque) to prevent batch effects from confounding differences between species. Samples were sequenced on an Illumina Hi-Seq 2000 or NextSeq500 at the Cornell University Biotechnology Resource Center.

#### Mapping PRO-seq reads

We mapped PRO-seq reads using standard informatics tools. Our PRO-seq mapping pipeline begins by removing reads that fail Illumina quality filters and trimming adapters using cutadapt with a 10% error rate^83^. Reads were mapped with BWA^84^ to the appropriate reference genome (either hg19, panTro4, rheMac3, mm10, or rn6) and a single copy of the Pol I ribosomal RNA transcription unit (GenBank ID# U13369.1). Mapped reads were converted to bigWig format for analysis using BedTools^85^ and the bedGraphToBigWig program in the Kent Source software package^86^. The location of the RNA polymerase active site was represented by the single base, the 3’ end of the nascent RNA, which is the position on the 5’ end of each sequenced read. After mapping reads to the reference genome, three samples (one human, U and PI, one chimpanzee, U and PI, and one rhesus macaque, U and PI) were identified as having poor data quality on the basis of the number of uniquely mapped reads, and were excluded from downstream analysis.

### Mapping 1:1 orthologs between different species

During all comparative analyses, the genomic coordinates of mapped reads, dREG scores, and other parameters of interest were converted to the human assembly (hg19) using CrossMap^87^. We converted genomic coordinates between genome assemblies using reciprocal-best (rbest) nets^88^. Reciprocal-best nets have the advantage that comparisons between species are constrained to 1:1 orthologues. This constraint on mapping is enforced by requiring each position to map uniquely in a reciprocal alignment between the human reference and the other species in the comparison. We downloaded rbest nets for hg19-mm10, hg19-panTro4, hg19-rn6 from the UCSC Genome Browser. We created rbest nets for hg19-rheMac3 using the doRecipBets.pl script provided as part of the Kent Source software package.

### Analysis of transcriptional regulatory elements

#### Defining a consensus set of transcriptional regulatory elements

We predicted TREs using dREG^54^ separately in each species’ reference genome. dREG uses a support vector regression model to score each site covered in a PRO-seq dataset based on its resemblance to features associated with transcription start sites in a reference training dataset. The dREG model was trained to recognize DNase-I-hypersensitive sites that also show substantial evidence of GRO-cap data in six PRO-seq or GRO-seq datasets measuring transcription in resting K562 cells. dREG scores were computed in the reference genome of each species in order to provide as much information as possible on the native context of each locus. In all cases, we combined the reads from all individuals for each species in order to maximize power for the discovery of TREs. In the primate species, treated and untreated CD4+ T-cells were analyzed separately.

We then defined a consensus set of TRE annotations, each of which bore the signature of an active TRE in at least one species and treatment condition. To define such a set, dREG scores were first converted to human reference genome (hg19) coordinates using CrossMap and the reciprocal-best nets. The advantage of converting dREG scores between the reference genome is that individual bases transfer more completely than genomic intervals using CrossMap and related tools. We then identified TREs in each species separately by thresholding the dREG scores. In all analyses, we selected a threshold of 0.3, which corresponds to a predicted false discovery rate of <7% compared with other sources of genomic data in human CD4+ T-cells. In addition, parallel analyses at separate thresholds (0.25 and 0.35) provided results that were in all cases consistent with those reported in the main manuscript (**Supplementary Table 2**). The set of overlapping TREs from each species were reduced to a single element containing the union of all positions covered by the set using bedops, and sites within 500 bp of each other were further merged. We assigned each putative TRE the maximum dREG score for each species and for each treatment condition.

#### Identifying differences in TREs between species

Differences in TRE transcription in 3-way (human-chimp-rhesus macaque) or 5-way (human-chimp-rhesus macaque-mouse-rat) species comparisons were identified using a combination of heuristics and statistical tests. Starting with the consensus set of TREs in hg19 coordinates, we first excluded potential one-to-many orthologs, by eliminating TREs that overlapped gaps in the reciprocal-best nets that were not classified as gaps in the standard nets. The remaining TREs were classified as unmappable when no orthologous position was defined in the rbest nets. Complete gains and losses were defined as TREs that were mappable in all species and for which the dREG score was less than 0.05 in at least one species and greater than 0.30 in at least one other species (see **Supplementary Note 1**). Gains and losses were assigned to a lineage based on an assumption of maximum parsimony under the known species phylogeny. We defined a set of TREs that displayed high-confidence changes in activity by comparing differences in PRO-seq read counts between species using deSeq2 and thresholding at a 1% false discovery rate (as described below). Changes in TRE activities were compared to histone modification ChIP-seq, DNase-I-seq, and DNA methyl immunoprecipitation data from the Epigenome Roadmap project^58^.

#### TRE classification

For some analyses, TREs were classified as likely promoters or enhancers on the basis of their distance from known protein-coding gene annotations (GENCODE v.19). TRE classes of primary interest include (see also **Supplementary Fig. 7**): (1) Promoters: near an annotated transcription start site (<100 bp); (2) Enhancers: distal to an annotated transcription start site (>5,000 bp)

#### Covariates that correlate with TRE changes

We compared the frequency at which evolutionary changes in transcription occur at TREs in a variety of different genomic contexts. We examined the rate of change as a function of distance from the nearest annotated transcription start site in GenCode v.19. TREs were binned by distance in increments of 0.02 on a log10 scale and we evaluated the mean rate at which evolutionary changes in TRE transcription arise in each bin. We also compared the rate of changes in TRE transcription across a variety of functional associations, including loop interactions, within the same topological associated domain, and in super-enhancers. H3K4me2 ChIA-PET data describing loop interactions were downloaded from the Gene Expression Omnibus (GEO) database (GSE32677) and the genomic locations of loops were converted from hg18 to hg19 coordinates using the liftOver tool. We also analyzed a separate dataset profiling loop interactions based on promoter capture Hi-C data in human CD4+ T-cells taken from the supplementary materials of ref. ^74^. Topological associated domains (TADs) based on Hi-C data for GM12878 cells were also downloaded from GEO (GSE63525). Super-enhancers in CD4+ T-cells were taken from the supplementary data for ref. ^75^. In all cases we excluded sites with potential one-to-many orthology in any of the species included in the comparison (typically just the three primates). Potential one-to-many orthologs were defined based on differences in the standard and reciprocal-best nets for each species pair.

#### Refining the location of active TREs using dREG-HD

During analyses of transcription factor binding motifs we further refined the location of TREs to the region between divergent paused RNA polymerase using a strategy that we call dREG-HD (manuscript in preparation, preliminary version available at https://github.com/Danko-Lab/dREG.HD). Briefly, we used an epsilon-support vector regression (SVR) with a Gaussian kernel to map the distribution of PRO-seq reads to smoothed DNase-I signal intensities. Training was conducted on randomly chosen positions within dREG peaks extended by 200bp on either side. Selection of feature vectors was optimized based on Pearson correlation coefficients between the imputed and experimental DNase-I score over the validation set. PRO-seq data was normalized by sequencing depth and further scaled such that the maximum value of any prediction dataset is within 90 percentile of the training examples. We chose a step size to be 60bp and extending 30 steps on each direction. The final model was trained using matched DNase-I and PRO-seq data in K562 cells.

Next we identified peaks in the imputed DNase-I hypersensitivity profile by fitting the imputed DNase-I signal using a cubic spline and identifying local maxima. We optimized two free parameters that control the (1) smoothness of spline curve fitting, and (2) threshold on the imputed DNase-I signal intensity. Parameters were optimized to achieve an appropriate trade-off between FDR and sensitivity on the testing K562 dataset. Parameters were tuned using a grid optimization over free parameters.

### DNA sequence analysis

#### Finding candidate transcription factor binding motifs

All motif analyses focused on 1,964 human TF binding motifs from the CisBP database^62^ clustered using an affinity propagation algorithm into 567 maximally distinct DNA binding specificities (see ref ^63^). Scores, which reflect a log_e_-odds ratio comparing each candidate motif model to a third-order Markov background model, were calculated using the RTFBSDB package^63^. We selected two separate motif thresholds for different analyses. Scores >10 were used in analyses which mix multiple TF binding motifs, and strike a tradeoff that focuses on minimizing false positives at the expense of sensitivity. We dropped the cutoff score to motifs >8 in analyses that use individual motifs in order to increase statistical power. For each of these thresholds, we estimated the mean genome-wide positive predictive values to be 0.47 and 0.38, respectively, for motif cutoffs of 10 and 8, by comparing motifs to ChIP-seq peak calls in K562 cells. During comparative analyses we scanned each primate reference genome separately with each motif to allow the detection of a putative binding site in any of the species included in the analysis, and then moved scores to a human (hg19) reference genome using the CrossMap tool. We chose this strategy because changes in TRE activity may reflect changes in binding in any of the primate species. For example, human gains may be explained by either a new binding site for a transcriptional activator in the human genome, or a loss in binding of a transcriptional repressor that was bound in both primate species.

#### Motif enrichment in TREs that change during CD4+ T-cell activation

Motifs enriched in up- or down-regulated dREG-HD TREs during CD4+ T-cell activation (*p* < 0.01) were selected using Fisher’s exact test with a Bonferroni correction for multiple hypothesis testing. Up- or down-regulated TREs were compared to a background set of >2,500 GC-content matched TREs that do not change transcription levels following π treatment (log-2 fold change <0.5-fold in magnitude and *p* > 0.25) using the *enrichmentTest* function in RTFBSDB^63^. To test for motif robustness, the background resampling was repeated 100 times and motifs were selected that achieve a significant result in >90%.

#### DNA sequence conservation analysis

For our evolutionary conservation analysis, we used phyloP scores^61^ based on the 100-way genome alignments available in the UCSC Genome Browser (hg19). In all cases, bigWig files were obtained from the UCSC Genome Browser and processed using the bigWig package in R. We represented evolutionary conservation as the mean phyloP score in each identified TFBS in the indicated set of dREG-HD sites.

#### Enrichment of DNA sequence changes in motifs

We identified single-nucleotide DNA sequence differences at sites at which two of three primate species share one base and the third species diverges. We intersected these species-specific divergences with matches to transcription factor binding motifs found within dREG-HD sites that undergo transcriptional changes between primate species. Because many motifs in Cis-BP are similar to one another, we first partitioned the motifs using clustering (as described above), and examined enrichments at the level of these clusters. Motifs were ranked by the Fisher’s exact test p-value of the enrichment of species divergences in dREG-HD sites that change transcription status (where changes in DNA sequence and transcription occur on the same branch) to dREG-HD sites that do not change. We also compute the enrichment ratio, which we define as the number of species divergences in each TF binding motif in dREG-HD sites that change on the same branch normalized to the same statistic in sites that do not change.

#### INSIGHT analysis

We examined the modes by which DNA sequences evolve in human lineage-specific dREG-HD sites or DHSs using INSIGHT^69^. We passed INSIGHT either complete DHSs, dREG-HD sites, or TFBS within dREG-HD sites that undergo the changes (see *Identifying differences in TREs between species*) indicated in the comparison. Human gains and losses, for example, were comprised of 4,384 dREG-HD sites with 9,924 separate regions (median length of 16 bp) after merging overlapping TFBSs with a log-odds score greater than 10. We also analyzed 24 transcription factors each of which has more than 900 occurrences in dREG-HD sites that change on the human branch (log-odds score >8). All analyses were conducted using the INSIGHT web server (http://compgen.cshl.edu/INSIGHT/) with the default settings enabled.

#### bQTL analysis

Frequency shift estimates for all variants in Teranchi et al. (2016) were provided by the Frasier lab and converted to a queryable database filtered to include only variants with coverage by 25 reads (75^th^ percentile) or more to avoid noise at low read counts. For each sequence/variant query, a set of four equivalent sequences/alternate allele pairs was constructed by swapping which allele was the reference and getting the reverse complement for both alleles. For example, given a sequence:variant:position combination of AATCGAA:C:3, the other queries produced were AACCGAA:T:3 (allele swap), TTCGATT:G:5 (reverse complement), and TTCGGTT:A:5 (reverse complement allele swap). Frequency shifts were computed by taking the post-ChIP frequency minus the pre-ChIP frequency for the human reference allele. Since k-mers longer than 7 had few hits, we allowed for wildcards (N) in longer sequences that would match any base. Wildcards were introduced into a k-mer by matching the k-mer sequence to the NF-kB motif and replacing the 3 lowest information content positions with N(s). Systematic shifts from 0 were tested using a one-tailed t-test. P-values for systematic differences at multiple sites were combined using Fisher’s method.

### De novo discovery of transcription units

#### Identification of transcription units (TU) using a three-state hidden Markov model

We inferred transcription units (TU) using a three-state hidden Markov model (HMM) similar to those we have recently published^51,70^. Each TU begins at a TRE identified using dREG and continues through the entire region inferred to be transcribed, which can covers tens- to hundreds- of kilobases. Three states were used to represent background (i.e., outside of a transcription unit), the TU body, and a post-polyA decay region. The HMM transition structure is shown in **Supplementary Fig. 13a**. We allow skipping over the post-polyA state, as unstable transcripts do not have these two-phase profiles. We took advantage of dREG as a potential signal for transcription initiation by incorporating the dREG score (maximum value in the interval from a given positive read-count position until the next, clamped to the zero-one interval) as a transition probability from the background to the transcription body state. PRO-seq data is generally sparse, so we applied a transformation that encoded only non-zero positions and the distance between such consecutive positions (**Supplementary Fig. 13a**). Our model described this transformed data using emissions distributions based on two types of variables. The first type of emission variable defines the PRO-seq read counts in non-zero positions. These counts were modeled using Poisson distributions in the background and post-polyA states, and using a Negative Binomial distribution in the transcription body state. The negative binomial distribution can be seen as a mixture of Poisson distributions with gamma-distributed rates and therefore allows for variation in TU expression levels across the genome. The second type of emission variable describes the distribution of distances in base pairs between positions having non-zero read counts. This distribution was modeled using a separate geometric distribution for each of the three states. Maximum likelihood estimates of all free parameters were obtained via Expectation Maximization, on a per-chromosome basis. TU predictions were then obtained using the Viterbi algorithm with parameters fixed at their maximum-likelihood values. Finally these predictions were mapped from the transformed coordinates back to genomic coordinates. Source code for our implementation is publicly available on GitHub: https://github.com/andrelmartins/tunits.nhp.

#### Inferring TU boundaries in the common great ape ancestor

We identified the most likely TU locations in the great ape ancestor by maximum parsimony. TUs were identified and compared in human reference coordinates (hg19) for all species. We used the bedops package to find the intersection between the predicted TU intervals in each pair of species (i.e., human-chimp, human-rhesus macaque, and chimp-rhesus macaque). Intersections (>= 1bp) between pairs of species were merged, resulting in a collection of TUs shared by any two pairs of species, and therefore likely to be a TU in the human-chimp ancestor. All steps were applied independently on the plus and minus strands. These steps identified 37,626 putative TUs active in CD4+ T-cells of the primate ancestor. We added 17,167 TUs that did not overlap ancestral TUs but were found in any one of the three primate species.

#### Transcription unit classification

TUs were classified by annotation type using a pipeline similar to ones that we have described recently^51,70,89^. Before classifying TUs we applied a heuristic to refine TUs on the basis of known annotations. TUs that completely overlap multiple gene annotations were broken at the transcription start site provided that a dREG site overlapped that transcription start site. Classification was completed using a set of rules to iteratively refine existing annotations, as shown in **Supplementary Fig. 13A**. Unless otherwise stated, overlap between a TU and a transcript annotation was defined such that >50% of a TU matched a gene annotation and covers at least 50% of the same annotation. TUs overlapping GENCODE annotations (>50% overlap, defined as above) were classified using the biotype in the GENCODE database into protein coding, lincRNA (lincRNA or processed transcript), or pseudogene. The remaining transcripts were classified as annotated RNA genes using GENCODE annotations, the rnaGenes UCSC Genome Browser track (converted from hg18 to hg19 coordinates), and miRBase v20^90^. As many RNA genes are processed from much longer TUs, we required no specific degree of overlap for RNA genes. Upstream antisense (i.e., divergent) TUs were classified as those within 500bp of the transcription start site of any GENCODE or higher level TU annotation (including lincRNAs). Antisense transcripts were defined as those with a high degree of overlap (>50%) with annotated protein coding genes in the opposite orientation. The remaining transcripts with a high degree of overlap (>50%) to annotated repeats in the repeatmasker database (rmsk) were classified as repeat transcription. Finally, any TUs still remaining were classified as unannotated, and were further divided into those which are intergenic or that partially overlapping existing annotations.

### Comparing transcription between conditions and species

#### Comparing transcription before and after CD4+ T-cell activation

We compared π treated and untreated CD4+ T-cells within each of the primate species using gene annotations (GENCODE v19). We counted reads in the interval between 500 bp downstream of the annotated transcription start site and either the end of the gene or 60,000 bp into the gene body (whichever was shorter). This window was selected to avoid (1) counting reads in the pause peak near the transcription start site, and (2) to focus on the 5’ end of the gene body affected by changes in transcription during 30 minutes of π treatment assuming a median elongation rate of 2 kb/ minute^51,91^. We limited analyses to gene annotations longer than 500 bp in length. To quantify transcription at enhancers, we counted reads in the window covered by each dREG-HD site plus an additional 250 bp on each end. Differential expression analysis was conducted using deSeq2^57^.

#### Comparing transcription between species

Read counts were compared between different species in hg19 coordinates. In all analyses, reads were transferred to the hg19 reference genome using CrossMap with rbest nets. Our analysis focused on transcription units or on the union of dREG sites across species. We focused our analysis of transcription units on the interval between 250 bp downstream of the annotated transcription start site and either the end of the gene or 60,000 bp into the gene body (whichever was shorter). We limited our analyses to TUs longer than 500 bp in length. Reads counts were obtained within each transcription unit, gene annotation, or enhancer, abbreviated here as a ‘region of interest’ (ROI), that has confident one-to-one orthology in all species examined in the analysis. This strategy of focusing on blocks of one-to-one orthology avoids errors caused by systematic differences in mappability or repeat content of species-specific genomic segments. We broke each each ROI into segments that have conserved orthology between hg19 and all species examined in the analysis, which included either a three-way (human-chimp-rhesus macaque) or five-way (human-chimp-rhesus macaque-mouse-rat) species comparison. We defined intervals of one-to-one orthology as those represented in levels 1, 3, and 5 of the reciprocal best nets (with gaps defined in levels 2, 4, and 6)^88^. Reads that map to regions that have orthology defined in all species were counted using the bigWig package in R using reads mapped to hg19 coordinates. Final counts for each ROI were defined as the sum of read counts within the regions of orthology that intersect that ROI. ROIs without confident one-to-one orthologs in all species analyzed were discarded. Our pipeline makes extensive use of the bigWig R package, Kent source tools, as well as the bedops and bedtools software packages^85,92^. Differential expression was conducted between species using the deSeq2 package for R, as described above.

### MCF-7 G11 cell culture

MCF7 G11 tamoxifen resistant cells, were a gift from Dr. Joshua LaBaer. Cells were maintained in DMEM with 5% FBS, antibiotics, and 1uM tamoxifen. MCF-7 G11 dCas9-KRAB stable cell lines were made (as described below) and were maintained in DMEM with 5% FBS, antibiotics, and 1uM tamoxifen. MCF-7 G11 dCas9-KRAB sgRNA stable cell lines were maintained in DMEM with 5% FBS, antibiotics, 1uM tamoxifen, and 150ug/ul Hygromycin B.

### Luciferase assays

Genomic DNA was isolated from human, chimp, and rhesus macaque PBMCs depleted for CD4+ cells using a Quick-DNA Miniprep Plus Kit (#D4068S; Zymo research) following the manufacturer’s instructions. Putative enhancer regions were amplified from the genomic DNA, restriction digested with KpnI and MluI, and cloned into the pGL3-promoter vector (Promega). The same orthologous regions were amplified from all three species with identical primers where possible or species-specific primers covering orthologous DNA in diverged regions. Vectors were co-transfected with pRL-SV40 Renilla (Promega) in a 10:1 ratio (500ng pGL3 to 50ng pRL-SV40) in MCF7 G11 cells cultured in 1uM tamoxifen. Transfected cells were treated with either 25ng/ml TNFa or water 21 hours after transfection. 24 hours post-transfection, luminescence was measured in triplicate using the Dual-Luciferase^®^ Reporter Assay System (Promega).

### Silencing endogenous TREs using dCAS9-KRAB

#### Cloning signle-guide RNAs (sgRNAs)

Single-guide RNAs (sgRNAs) were designed using the CRISPR design tool (http://crispr.mit.edu) and sequences are shown in **Supplementary Table 3**. Forward and reverse sgRNAs were synthesized separately by IDT and annealed. T4 Polynucleotide Kinase (NEB) was used to phosphorylate the forward and reverse sgRNA during the annealing. 10x T4 DNA Ligase Buffer, which contains 1mM ATP, was incubated for 30 minutes at 37°C and then at 95C for 5 minutes, decreasing by 5°C every 1 minute until 25°C. Oligos were diluted 1:200 using Molecular grade water. sgRNAs were inserted into the pLenti SpBsmBI sgRNA Hygro plasmid from addgene (#62205) by following the authors protocol^93^. The plasmid was linearized using BsmBI digestion (NEB) and purified using gel extraction (QIAquick Gel Extraction Kit). The purified linear plasmid was then dephosphorylated using Alkaline Phosphatase Calf Intestinal (CIP) (NEB) to ensure the linear plasmid did not ligate with itself. A second gel extraction was used as before to purify the linearized plasmid. The purified dephosphorylated linear plasmid and phosphorylated annealed oligos were ligated together using the Quick Ligation Kit (NEB). The ligated product was transformed into One Shot Stbl3 Chemically Competent E. coli (ThermoFisher Scientific). 100ul of the transformed bacteria were plated on Ampicillin (200ug/ml) plates. Single colonies were picked, sequenced, and the plasmid was isolated using endo free midi-preps from Omega.

#### Transfection of MCF-7 G11 cell lines

We used lentivirus to transfect MCF-7 cells. Lentivirus was made using lipofectamine 3000 from Invitrogen. Phoenix Hek cells (grown in DMEM with 10% FBS and antibiotics) were seeded in a 6-well plate at 400,000 cells/plate. Cells were grown until ∼90% confluent. 1ug of pHAGE_EF1a_dCas9-KRAB plasmid from addgene (#50919) plasmid or the pLenti SpBsmBI sgRNA Hygro (addgene #62205) containing each sgRNA, 0.5ug of psPAX (addgene #12260), and 0.25ug pMD2.G (addgene #12259) were mixed.

MCF-7 G11 cells were plated at ∼200,000 cells/well in a 6-well plate. 24 hours later 3ml/well of virus was mixed with 10ug/ml polybrene and incubated for 5 minutes at room temperature. This mix was added to the cells and centrifuged for 40 minutes at 800g at 32C (Viju Vijayan Pillai, personal communication). 12-24 hours later the virus was removed and fresh media was added. 24-48 hours later the cells were selected with 2ug/ml puromycin for 2 weeks. The MCF-7 G11 dCas9-KRAB stable cell lines was grown and maintained in puromycin. A second lentiviral infection was done using the stable MCF-7 G11 dCas9-KRAB cells. The same protocol was used. 24-48 hours later the cells were selected with 150ug/ml Hygromycin B. New stable cell lines were grown and maintained in hygromycin B.

#### TNFa treatment

Prior to TNFa treatment, cells were grown for 3 days in DMEM with 5% FBS, antibiotics, tamoxifen and hygromycin. Cells were then left untreated or treated for 40 min with 25ng/ml TNFa. RNA was extracted using TRIzol Reagent (Invitrogen). We reverse transcribed 1ug of RNA and used this as input for real-time quantitative PCR (RT-PCR) to analyze *SGPP2* expression. Primers for *SGPP2* were designed targeting a sequence in intron 1, upstream of the intronic enhancer. Raw *Cp* values were transferred to units of expression using a standard dilution curve comprised of a mixture of cDNA from each sample within the biological replicate. We included four serial dilutions, each of which covered a two-fold difference in expression. Each sample was further normalized for differences in RNA content by primers recognizing the 18S rRNA control. The ratio between normalized *SGPP2* expression in each sgRNA-transfected MCF-7 cell line and the empty vector control was log-2 transformed and tested for differences from 0 using a two-sided t-test.

### Data availability

PRO-seq data was deposited into the Gene Expression Omnibus database under accession number GSE85337.

### Code availability

All data analysis scripts and software are publicly available on GitHub: https://github.com/Danko-Lab/CD4-Cell-Evolution.

## Acknowledgements

We thank M. Jin for assistance in establishing the magnetic separation of CD4+ T-cells, L. Core, H. Kwak, N. Fuda, and I. Jonkers for assistance troubleshooting the PRO-seq library prep, and A. Wetterau for preparing nuclei for mouse and rat CD4+ T-cells. Work in this publication was supported by generous seed grants from the Cornell University Center for Vertebrate Genomics (CVG), the Center for Comparative and Population Genetics (3CPG), NHGRI (National Human Genome Research Institute) grant HG009309 to CGD, NHLBI (National Heart, Lung, and Blood Institute) grant UHL129958A to CGD and JTL, NIGMS (National Institute of General Medical Sciences) grant GM102192 to AS, NHGRI (National Human Genome Research Institute) grant HG0070707 to AS and JTL, NIH/NIDDK DK058110 to WLK, and CPRIT RP160319 to WLK. The content is solely the responsibility of the authors and does not necessarily represent the official views of the US National Institutes of Health. Finally, we would like to thank the anonymous human and non-human primate donors who gave blood in support of this study.

## Author contributions

LAC, BAM, CGD, EJR, and ETW performed CD4+ T-cell extraction, validation, and PRO-seq experiments. CGD, ZW, TC, ALM, LAC, and ND analyzed the data. CGD, AS, JTL, WLK, and SAC supervised data collection and analysis. CGD and AS wrote the paper with input from the other authors.

## Competing financial interests

The authors declare no competing financial interests.

## Author information

PRO-seq data was deposited into the Gene Expression Omnibus database under accession number GSE85337. All data analysis scripts and software are publicly available on GitHub: https://github.com/Danko-Lab/CD4-Cell-Evolution.

## Supplementary Figure Legends

**Supplementary Fig. 1 | Validation of CD4+ cell enrichment by flow cytometry.** Representative plots of CD4 expression in human, chimpanzee, and rhesus macaque PBMC, before (left) and after (right) CD4 microbead enrichment. Percentage of total live lymphocytes shown.

**Supplementary Fig. 2 | Transcription abundance in the gene bodies of T-cell lineage specific markers.** Plots show normalized expression (log2 scale) of transcription factors and cytokines that mark specific subsets of CD4+ T-cell population in the species indicated below the plot. Each point represents the transcription of the indicated gene in a different untreated T-cell sample. The bar indicates the mean in each species. In all cases read counts were limited to regions of orthology in the bodies of genes indicated on each plot.

**Supplementary Fig. 3 | Principal component analysis (PCA) of CD4+ T-cell PRO-seq libraries.** Scatterplots show the first five principal components (PC) from CD4+ T-cell PRO-seq libraries. PCA was constructed using regions of orthology in all five species in the bodies of transcription units identified by a three state hidden Markov model. The fraction of the variance explained by each PC is shown in parentheses. The key shown below the plot indicates the species and treatment condition of each point.

**Supplementary Fig. 4 | Sample correlation plotted as a function of estimated evolutionary divergence time between species.** Scatterplot shows the evolutionary divergence time (X-axis) as a function of Spearman’s correlation in gene body transcription between each sample collected in the untreated condition and the mean gene expression in untreated human CD4+ T-cells (Y-axis). The red line shows the best linear fit and dotted lines indicate the 99% confidence interval. We assume the following evolutionary divergence estimates for each species pair with respect to human, 12 MYR for chimp-human [Moorjani et. al. (2016)], 25 MYR for human-rhesus [Rogers (2013)], and 75 MYR for human-rodent [Chinwalla et. al. (2002)].

**Supplementary Fig. 5 | Changes in gene transcription following PMA+Ionomycin treatment in chimpanzee and rhesus macaque CD4+ T-cells. (a-b)** MA plot shows the log-2 fold change following π treatment (y-axis) as a function of the mean transcription level in GENCODE annotated genes (x-axis) in data from chimpanzee (left) and rhesus macaque (right) CD4+ T-cells. Red points indicate statistically significant changes (p < 0.01). Several classical response genes that undergo well-documented changes in transcript abundance following CD4+ T-cell activation (e.g., *IL2*, *IFNG*, *TNF*, and *EGR3*) are marked. **(c)** UCSC genome browser track shows transcription in the *IFNG* locus in untreated (U) and PMA+ionomycin (π) treated CD4+ T-cells isolated from the primate species indicated at left. PRO-seq tracks show transcription on the plus (red) and minus (blue) strands. dREG tracks show the distribution of dREG signal. The net-synteny tracks show the fraction of the genomic area that is mappable in the indicated species. The location of transcription units inferred in the common ancestor of human and chimpanzee, and the location of RefSeq gene annotations, are shown at the bottom. **(d-f)** Scatterplots show the correlation between changes in gene expression (log-2 scale) following π treatment in the species indicated on the axes. Color scale indicates the density of points in the region.

**Supplementary Fig. 6 | Evolutionary changes in TREs. (a)** Venn diagram illustrating raw changes in TREs among primate species. In all cases, TREs were discovered in untreated CD4+ T-cells using dREG (threshold > 0.3). **(b)** Q-Q plot showing observed p-values (deSeq2 in human compared to the other two primate species) among TREs that were not identified by dREG in at least one species (red), all TREs identified (black), and a set of conserved TREs (gray).

**Supplementary Fig. 7 | Evolutionary changes in TREs correlate with chromatin and DNA modifications.** ChIP-seq signal for H3K27ac and H3K4me1 near dREG sites classified as gains, losses, or complete losses of TRE signal (dREG score < 0.05) on the human branch.

**Supplementary Fig. 8 | PhyloP scores in transcription factor (TF) binding motifs. (a)** Evolutionary conservation centered on matches to a TF binding motif at the indicated cut off score (left), or adjusted for distance to the nearest annotated transcription start site by subsampling (right) **(b)** PhyloP scores that fall within the binding motifs recognized by STAT2 (M6494_1.02), YY1 (M4490_1.02), CREB1 (M6180_1.02), and ELF1 (M6203_1.02). In all cases motifs fall in dREG-HD that are gained (blue) or lost (cyan) on the human branch, or are conserved among all primate species (red). **(c)** The distribution of human derived alleles near dREG sites that are gained (blue) or lost (cyan) on the human branch, or are conserved among all primate species (red).

**Supplementary Fig. 9 | Candidate causal DNA sequence differences underlying changes in *SGPP2* transcription.** UCSC genome browser track shows transcription near *SGPP2* and *FARSB* in untreated (U) and PMA+ionomycin (π) treated human CD4+ T-cells or in human MCF-7 cells. PRO-seq tracks show transcription on the plus (red) and minus (blue) strands. Axes for the PRO-seq data are in units of reads per kilobase per million mapped (RPKM) or in raw reads (MCF-7). dREG tracks show the distribution of dREG signal. Heatmap (top) shows Hi-C signal in GM12878 lymphoblastoid cell lines. Insert (bottom) shows lack of orthology in chimpanzee and rhesus macaque in an active TRE (human) that binds a number of TFs in ENCODE cell lines (left) and substitutions in NF-kB binding motifs near *SGPP2*. Two motif occurrences in the proximal promoter were bound by RELA, a subunit of NF-kB, based on human ChIP-seq data in ENCODE cell lines (green boxes). Positions where human carries a derived allele are indicated by yellow highlights. PRO-seq reads matched the human reference allele in all positions (15/ 15 reads match C and 26/ 26 match the reference T allele in the NF-kB binding site in the promoter; 11/ 11 reads match the G and 11/ 11 match the T reference allele in the NF-kB binding site in the promoter; and 24/ 24 reads support the TG human reference sequence in the internal enhancer). Scatterplots show the relative frequencies of the human allele in RELA (NF-kB) ChIP-seq data matching NF-kB binding QTLs that mimic the human and ancestral alleles, while controlling for the flanking sequence indicated below the plot. The red dot denotes the mean. All four human-specific DNA sequence changes in NF-kB binding motifs in the proximal promoter together show trend toward higher NF-kB binding in human (*p* = 0.017, using Fisher’s method to combine p-values).

**Supplementary Fig. 10 | Luciferase assays for TREs identified near *SGPP2.*** The Y-axis shows the luciferase signal driven by the *SGPP2* promoter or the internal enhancer in MCF-7 cells using DNA from each primate species following 3 hours of stimulation with TNFα or vehicle control. Bars show the mean luciferase activity in each species, over the empty vector and renilla controls. Error bars represent the standard error of the mean.

**Supplementary Fig. 11 | Adaptive substitutions in specific TF binding motifs.** Adaptive substations in TF binding motifs (TFBM) occurring commonly (>900 times) in human lineage-specific dREG-HD sites. Columns denote the TF name annotated in CisBP (TF), number of sites (Sites), the number of bases (Bases), the expected number of adaptive substitutes per kilobase (E[A]), the standard error in the expected substitutions per kilobase (E[A]_stderr), and the estimated number of adaptive substitutions (# Adaptive Substitutions). TFBSs may be bound by any TF that recognizes a similar motif. TFBM in which E[A] is significantly larger than 0 are highlighted in bold fold. The estimated number of adaptive substitutions for each of these sites is shown.

**Supplementary Fig. 12 | Discovery of transcription units (TU) in primate T-cells. (a)** A novel three-state hidden Markov model (HMM) was used to discover transcription units. States correspond to non-transcribed background sequence, transcribed sequence, and post polyA transcription. TUs were classified into one of seven classes as indicated in the cartoon. **(b)** The number and fraction of transcription units that fall into each TU classification. **(c)** Example of the hidden Markov model (HMM) in a typical region. TUs largely agree with RefSeq gene annotations when available.

**Supplementary Fig. 13 | DNA sequence conservation as a function of genomic distance to the nearest start site.** Scatterplot shows the percentage of TREs undergoing complete gains and losses (left), undergoing a partial change in the abundance of Pol II (center), or that are not alignable between species (right) as a function of distance from the nearest annotated transcription start site (x-axis). The size of each point represents the amount of data in the corresponding distance bin.

**Supplementary Fig. 14 | Evolutionary conservation of DNA sequence mirrors functional conservation at looped- and un-looped enhancers.** Cumulative distribution function of phyloP scores from the 100-way alignments in the indicated class of dREG site. The insert shows the fraction of sites in each class exceeding a phyloP score cutoff of 0.75.

## References

1. Jacob, F. & Monod, J. Genetic regulatory mechanisms in the synthesis of proteins. J. Mol. Biol. 3, 318–356 (1961).

2. Britten, R. J. & Davidson, E. H. Gene regulation for higher cells: a theory. Science 165, 349–357 (1969).

3. King, M. C. & Wilson, A. C. Evolution at two levels in humans and chimpanzees. Science 188, 107–116 (1975).

4. Rockman, M. V. et al. Ancient and recent positive selection transformed opioid cis-regulation in humans. PLoS Biol. 3, e387 (2005).

5. Prabhakar, S. et al. Human-specific gain of function in a developmental enhancer. Science 321, 1346–1350 (2008).

6. Capra, J. A., Erwin, G. D., McKinsey, G., Rubenstein, J. L. R. & Pollard, K. S. Many human accelerated regions are developmental enhancers. Philos. Trans. R. Soc. Lond. B Biol. Sci. 368, 20130025 (2013).

7. McLean, C. Y. et al. Human-specific loss of regulatory DNA and the evolution of human-specific traits. Nature 471, 216–219 (2011).

8. Cotney, J. et al. The evolution of lineage-specific regulatory activities in the human embryonic limb. Cell 154, 185–196 (2013).

9. Arbiza, L. et al. Genome-wide inference of natural selection on human transcription factor binding sites. Nat. Genet. 45, 723–729 (2013).

10. Prescott, S. L. et al. Enhancer Divergence and cis-Regulatory Evolution in the Human and Chimp Neural Crest. Cell 163, 68–83 (2015).

11. Siepel, A. & Arbiza, L. Cis-regulatory elements and human evolution. Curr. Opin. Genet. Dev. 29, 81–89 (2014).

12. Wray, G. A. The evolutionary significance of cis-regulatory mutations. Nat. Rev. Genet. 8, 206–216 (2007).

13. Wilson, M. D. et al. Species-specific transcription in mice carrying human chromosome 21. Science 322, 434–438 (2008).

14. Khurana, E. et al. Integrative annotation of variants from 1092 humans: application to cancer genomics. Science 342, 1235587 (2013).

15. Haygood, R., Fedrigo, O., Hanson, B., Yokoyama, K.-D. & Wray, G. A. Promoter regions of many neural- and nutrition-related genes have experienced positive selection during human evolution. Nat. Genet. 39, 1140–1144 (2007).

16. Torgerson, D. G. et al. Evolutionary processes acting on candidate cis-regulatory regions in humans inferred from patterns of polymorphism and divergence. PLoS Genet. 5, e1000592 (2009).

17. Schmidt, D. et al. Five-vertebrate ChIP-seq reveals the evolutionary dynamics of transcription factor binding. Science 328, 1036–1040 (2010).

18. Ballester, B. et al. Multi-species, multi-transcription factor binding highlights conserved control of tissue-specific biological pathways. Elife 3, e02626 (2014).

19. Vierstra, J. et al. Mouse regulatory DNA landscapes reveal global principles of cis-regulatory evolution. Science 346, 1007–1012 (2014).

20. Arnold, C. D. et al. Quantitative genome-wide enhancer activity maps for five Drosophila species show functional enhancer conservation and turnover during cis-regulatory evolution. Nat. Genet. 46, 685–692 (2014).

21. Doniger, S. W. & Fay, J. C. Frequent gain and loss of functional transcription factor binding sites. PLoS Comput. Biol. 3, e99 (2007).

22. Zheng, W., Zhao, H., Mancera, E., Steinmetz, L. M. & Snyder, M. Genetic analysis of variation in transcription factor binding in yeast. Nature 464, 1187–1191 (2010).

23. Bradley, R. K. et al. Binding site turnover produces pervasive quantitative changes in transcription factor binding between closely related Drosophila species. PLoS Biol. 8, e1000343 (2010).

24. Villar, D. et al. Enhancer evolution across 20 mammalian species. Cell 160, 554–566 (2015).

25. Chuong, E. B., Elde, N. C. & Feschotte, C. Regulatory evolution of innate immunity through co-option of endogenous retroviruses. Science 351, 1083–1087 (2016).

26. Fuda, N. J., Ardehali, M. B. & Lis, J. T. Defining mechanisms that regulate RNA polymerase II transcription in vivo. Nature 461, 186–192 (2009).

27. Carvunis, A.-R. et al. Evidence for a common evolutionary rate in metazoan transcriptional networks. Elife 4, (2015).

28. Zhou, X. et al. Epigenetic modifications are associated with inter-species gene expression variation in primates. Genome Biol. 15, 547 (2014).

29. Cain, C. E., Blekhman, R., Marioni, J. C. & Gilad, Y. Gene expression differences among primates are associated with changes in a histone epigenetic modification. Genetics 187, 1225–1234 (2011).

30. Xiao, S. et al. Comparative epigenomic annotation of regulatory DNA. Cell 149, 1381–1392 (2012).

31. Paris, M. et al. Extensive divergence of transcription factor binding in Drosophila embryos with highly conserved gene expression. PLoS Genet. 9, e1003748 (2013).

32. Cusanovich, D. A., Pavlovic, B., Pritchard, J. K. & Gilad, Y. The functional consequences of variation in transcription factor binding. PLoS Genet. 10, e1004226 (2014).

33. Wong, E. S. et al. Decoupling of evolutionary changes in transcription factor binding and gene expression in mammals. Genome Res. 25, 167–178 (2015).

34. Hah, N., Murakami, S., Nagari, A., Danko, C. G. & Kraus, W. L. Enhancer transcripts mark active estrogen receptor binding sites. Genome Res. 23, 1210–1223 (2013).

35. Domené, S. et al. Enhancer turnover and conserved regulatory function in vertebrate evolution. Philos. Trans. R. Soc. Lond. B Biol. Sci. 368, 20130027 (2013).

36. Wunderlich, Z. et al. Krüppel Expression Levels Are Maintained through Compensatory Evolution of Shadow Enhancers. Cell Rep. 12, 1740–1747 (2015).

37. Cannavò, E. et al. Shadow Enhancers Are Pervasive Features of Developmental Regulatory Networks. Curr. Biol. 26, 38–51 (2016).

38. Ludwig, M. Z., Bergman, C., Patel, N. H. & Kreitman, M. Evidence for stabilizing selection in a eukaryotic enhancer element. Nature 403, 564–567 (2000).

39. Sanyal, A., Lajoie, B. R., Jain, G. & Dekker, J. The long-range interaction landscape of gene promoters. Nature 489, 109–113 (2012).

40. Vietri Rudan, M. et al. Comparative Hi-C reveals that CTCF underlies evolution of chromosomal domain architecture. Cell Rep. 10, 1297–1309 (2015).

41. Khan, Z. et al. Primate transcript and protein expression levels evolve under compensatory selection pressures. Science 342, 1100–1104 (2013).

42. Bauernfeind, A. L. et al. Evolutionary Divergence of Gene and Protein Expression in the Brains of Humans and Chimpanzees. Genome Biol. Evol. 7, 2276–2288 (2015).

43. Battle, A. et al. Genomic variation. Impact of regulatory variation from RNA to protein. Science 347, 664–667 (2015).

44. Pai, A. A. et al. The contribution of RNA decay quantitative trait loci to inter-individual variation in steady-state gene expression levels. PLoS Genet. 8, e1003000 (2012).

45. Core, L. J., Waterfall, J. J. & Lis, J. T. Nascent RNA sequencing reveals widespread pausing and divergent initiation at human promoters. Science 322, 1845–1848 (2008).

46. Kwak, H., Fuda, N. J., Core, L. J. & Lis, J. T. Precise maps of RNA polymerase reveal how promoters direct initiation and pausing. Science 339, 950–953 (2013).

47. Churchman, L. S. & Weissman, J. S. Nascent transcript sequencing visualizes transcription at nucleotide resolution. Nature 469, 368–373 (2011).

48. Nojima, T. et al. Mammalian NET-Seq Reveals Genome-wide Nascent Transcription Coupled to RNA Processing. Cell 161, 526–540 (2015).

49. Mahat, D. B. et al. Base-pair-resolution genome-wide mapping of active RNA polymerases using precision nuclear run-on (PRO-seq). Nat. Protoc. 11, 1455–1476 (2016).

50. Schwalb, B. et al. TT-seq maps the human transient transcriptome. Science 352, 1225–1228 (2016).

51. Hah, N. et al. A rapid, extensive, and transient transcriptional response to estrogen signaling in breast cancer cells. Cell 145, 622–634 (2011).

52. Core, L. J. et al. Analysis of nascent RNA identifies a unified architecture of initiation regions at mammalian promoters and enhancers. Nat. Genet. 46, 1311–1320 (2014).

53. Andersson, R. et al. Nuclear stability and transcriptional directionality separate functionally distinct RNA species. Nat. Commun. 5, 5336 (2014).

54. Danko, C. G. et al. Identification of active transcriptional regulatory elements from GRO-seq data. Nat. Methods 12, 433–438 (2015).

55. Khaitovich, P. et al. A Neutral Model of Transcriptome Evolution. PLoS Biol. 2, e132 (2004).

56. Dukler, N. et al. Nascent RNA sequencing reveals a dynamic global transcriptional response at genes and enhancers to the natural medicinal compound celastrol. bioRxiv 117689 (2017). doi:10.1101/117689

57. Love, M. I., Huber, W. & Anders, S. Moderated estimation of fold change and dispersion for RNA-seq data with DESeq2. Genome Biol. 15, 550 (2014).

58. Roadmap Epigenomics Consortium et al. Integrative analysis of 111 reference human epigenomes. Nature 518, 317–330 (2015).

59. Bonn, S. et al. Tissue-specific analysis of chromatin state identifies temporal signatures of enhancer activity during embryonic development. Nat. Genet. 44, 148–156 (2012).

60. Zentner, G. E., Tesar, P. J. & Scacheri, P. C. Epigenetic signatures distinguish multiple classes of enhancers with distinct cellular functions. Genome Res. 21, 1273–1283 (2011).

61. Pollard, K. S., Hubisz, M. J., Rosenbloom, K. R. & Siepel, A. Detection of nonneutral substitution rates on mammalian phylogenies. Genome Res. 20, 110–121 (2010).

62. Weirauch, M. T. et al. Determination and inference of eukaryotic transcription factor sequence specificity. Cell 158, 1431–1443 (2014).

63. Wang, Z., Martins, A. L. & Danko, C. G. RTFBSDB: an integrated framework for transcription factor binding site analysis. Bioinformatics (2016). doi:10.1093/bioinformatics/btw338

64. Macian, F. NFAT proteins: key regulators of T-cell development and function. Nat. Rev. Immunol. 5, 472–484 (2005).

65. Tehranchi, A. K. et al. Pooled ChIP-Seq Links Variation in Transcription Factor Binding to Complex Disease Risk. Cell 165, 730–741 (2016).

66. Franco, H. L., Nagari, A. & Kraus, W. L. TNFα signaling exposes latent estrogen receptor binding sites to alter the breast cancer cell transcriptome. Mol. Cell 58, 21–34 (2015).

67. Thakore, P. I. et al. Highly specific epigenome editing by CRISPR-Cas9 repressors for silencing of distal regulatory elements. Nat. Methods 12, 1143–1149 (2015).

68. Green, R. E. et al. A draft sequence of the Neandertal genome. Science 328, 710–722 (2010).

69. Gronau, I., Arbiza, L., Mohammed, J. & Siepel, A. Inference of natural selection from interspersed genomic elements based on polymorphism and divergence. Mol. Biol. Evol. 30, 1159–1171 (2013).

70. Chae, M., Danko, C. G. & Kraus, W. L. groHMM: a computational tool for identifying unannotated and cell type-specific transcription units from global run-on sequencing data. BMC Bioinformatics 16, 222 (2015).

71. Engreitz, J. M. et al. Local regulation of gene expression by lncRNA promoters, transcription and splicing. Nature 539, 452–455 (2016).

72. Chepelev, I., Wei, G., Wangsa, D., Tang, Q. & Zhao, K. Characterization of genome-wide enhancer-promoter interactions reveals co-expression of interacting genes and modes of higher order chromatin organization. Cell Res. 22, 490–503 (2012).

73. Rao, S. S. P. et al. A 3D map of the human genome at kilobase resolution reveals principles of chromatin looping. Cell 159, 1665–1680 (2014).

74. Javierre, B. M. et al. Lineage-Specific Genome Architecture Links Enhancers and Non-coding Disease Variants to Target Gene Promoters. Cell 167, 1369–1384.e19 (2016).

75. Hnisz, D. et al. Super-enhancers in the control of cell identity and disease. Cell 155, 934–947 (2013).

76. Barreiro, L. B., Marioni, J. C., Blekhman, R., Stephens, M. & Gilad, Y. Functional comparison of innate immune signaling pathways in primates. PLoS Genet. 6, e1001249 (2010).

77. Gilad, Y., Oshlack, A., Smyth, G. K., Speed, T. P. & White, K. P. Expression profiling in primates reveals a rapid evolution of human transcription factors. Nature 440, 242–245 (2006).

78. Blekhman, R., Oshlack, A., Chabot, A. E., Smyth, G. K. & Gilad, Y. Gene regulation in primates evolves under tissue-specific selection pressures. PLoS Genet. 4, e1000271 (2008).

79. Kutter, C. et al. Rapid turnover of long noncoding RNAs and the evolution of gene expression. PLoS Genet. 8, e1002841 (2012).

80. Lewis, J. J., van der Burg, K. R. L., Mazo-Vargas, A. & Reed, R. D. ChIP-Seq-Annotated Heliconius erato Genome Highlights Patterns of cis-Regulatory Evolution in Lepidoptera. Cell Rep. 16, 2855–2863 (2016).

81. Gilad, Y., Oshlack, A. & Rifkin, S. A. Natural selection on gene expression. Trends Genet. 22, 456–461 (2006).

82. Bailey, S. D. et al. Noncoding somatic and inherited single-nucleotide variants converge to promote ESR1 expression in breast cancer. Nat. Genet. 48, 1260–1266 (2016).

83. Martin, M. Cutadapt removes adapter sequences from high-throughput sequencing reads. EMBnet.journal 17, 10–12 (2011).

84. Li, H. & Durbin, R. Fast and accurate short read alignment with Burrows-Wheeler transform. Bioinformatics 25, 1754–1760 (2009).

85. Quinlan, A. R. & Hall, I. M. BEDTools: a flexible suite of utilities for comparing genomic features. Bioinformatics 26, 841–842 (2010).

86. Kuhn, R. M., Haussler, D. & Kent, W. J. The UCSC genome browser and associated tools. Brief. Bioinform. 14, 144–161 (2013).

87. Zhao, H. et al. CrossMap: a versatile tool for coordinate conversion between genome assemblies. Bioinformatics 30, 1006–1007 (2014).

88. Kent, W. J., Baertsch, R., Hinrichs, A., Miller, W. & Haussler, D. Evolution’s cauldron: duplication, deletion, and rearrangement in the mouse and human genomes. Proc. Natl. Acad. Sci. U. S. A. 100, 11484–11489 (2003).

89. Luo, X., Chae, M., Krishnakumar, R., Danko, C. G. & Kraus, W. L. Dynamic reorganization of the AC16 cardiomyocyte transcriptome in response to TNFα signaling revealed by integrated genomic analyses. BMC Genomics 15, 155 (2014).

90. Kozomara, A. & Griffiths-Jones, S. miRBase: annotating high confidence microRNAs using deep sequencing data. Nucleic Acids Res. 42, D68–73 (2014).

91. Danko, C. G. et al. Signaling pathways differentially affect RNA polymerase II initiation, pausing, and elongation rate in cells. Mol. Cell 50, 212–222 (2013).

92. Neph, S. et al. BEDOPS: high-performance genomic feature operations. Bioinformatics 28, 1919–1920 (2012).

93. Pham, H., Kearns, N. A. & Maehr, R. Transcriptional Regulation with CRISPR/Cas9 Effectors in Mammalian Cells. Methods Mol. Biol. 1358, 43–57 (2016).

